# Horizontal transfer of nuclear DNA in transmissible cancer

**DOI:** 10.1101/2024.07.26.604742

**Authors:** Kevin Gori, Adrian Baez-Ortega, Andrea Strakova, Maximilian R Stammnitz, Jinhong Wang, Jonathan Chan, Katherine Hughes, Sophia Belkhir, Maurine Hammel, Daniela Moralli, James Bancroft, Edward Drydale, Karen M Allum, María Verónica Brignone, Anne M Corrigan, Karina F de Castro, Edward M Donelan, Ibikunle A Faramade, Alison Hayes, Nataliia Ignatenko, Rockson Karmacharya, Debbie Koenig, Marta Lanza-Perea, Adriana M Lopez Quintana, Michael Meyer, Winifred Neunzig, Francisco Pedraza-Ordoñez, Yoenten Phuentshok, Karma Phuntsho, Juan C Ramirez-Ante, John F Reece, Sheila K Schmeling, Sanjay Singh, Lester J Tapia Martinez, Marian Taulescu, Samir Thapa, Sunil Thapa, Mirjam G van der Wel, Alvaro S Wehrle-Martinez, Michael R Stratton, Elizabeth P Murchison

## Abstract

Although somatic cell genomes are usually entirely clonally inherited, nuclear DNA exchange between cells of an organism can occur sporadically by cell fusion, phagocytosis or other mechanisms^1–3^. This phenomenon has long been noted in the context of cancer, where it could be envisaged that DNA horizontal transfer plays a functional role in disease evolution^4–13^. However, an understanding of the frequency and significance of this process in naturally occurring tumours is lacking. The host-tumour genetic discordance of transmissible cancers, malignant clones which pass between animals as allogeneic grafts, provides an opportunity to investigate this. We screened for host-to-tumour horizontal transfer of nuclear DNA in 174 tumours from three transmissible cancers affecting dogs and Tasmanian devils, and detected a single instance in the canine transmissible venereal tumour (CTVT). This involved introduction of a 15-megabase dicentric genetic element, composed of 11 rearranged fragments of six chromosomes, to a CTVT sublineage occurring in Asia around 2,000 years ago. The element forms the short arm of a small submetacentric chromosome, and derives from a dog with ancestry associated with the ancient Middle East. The introduced DNA fragment is transcriptionally active and has adopted the expression profile of CTVT. Its 143 genes do not, however, confer any obvious advantage to its spatially restricted CTVT sublineage. Our findings indicate that nuclear DNA horizontal transfer, although likely a rare event in tumour evolution, provides a viable mechanism for the acquisition of genetic material in naturally occurring cancer genomes.

## Main

Horizontal gene transfer is a potential source of adaptive variation in cancer cells^4–13^. Its detection, however, requires well-resolved cell phylogenies and sufficient genetic markers to distinguish donor and recipient DNA. This information is usually lacking in ordinary cancers which arise from cells of the genetically matched host. Transmissible cancers, however, malignant clones which pass between individuals as contagious allografts, are useful models with which to investigate this. The three naturally occurring transmissible cancers known in mammals are canine transmissible venereal tumour (CTVT), a sexually transmitted genital cancer affecting dogs; and two facial tumour clones, known as devil facial tumour 1 (DFT1) and devil facial tumour 2 (DFT2), which are spread by biting among Tasmanian devils^14^. Although transmissible cancers themselves arise rarely, once established in populations they can survive over long time periods^14–16^. Indeed, CTVT first emerged several thousand years ago and is today found in dogs worldwide (Figure 1A)^17–20^. Importantly, all tumours of a transmissible cancer carry the genome of the “founder animal” that spawned the lineage^21^. Variation in this germline constitutive genotype among tumours may indicate horizontal transfer of DNA from a transient allogeneic host.

**Figure 1:**
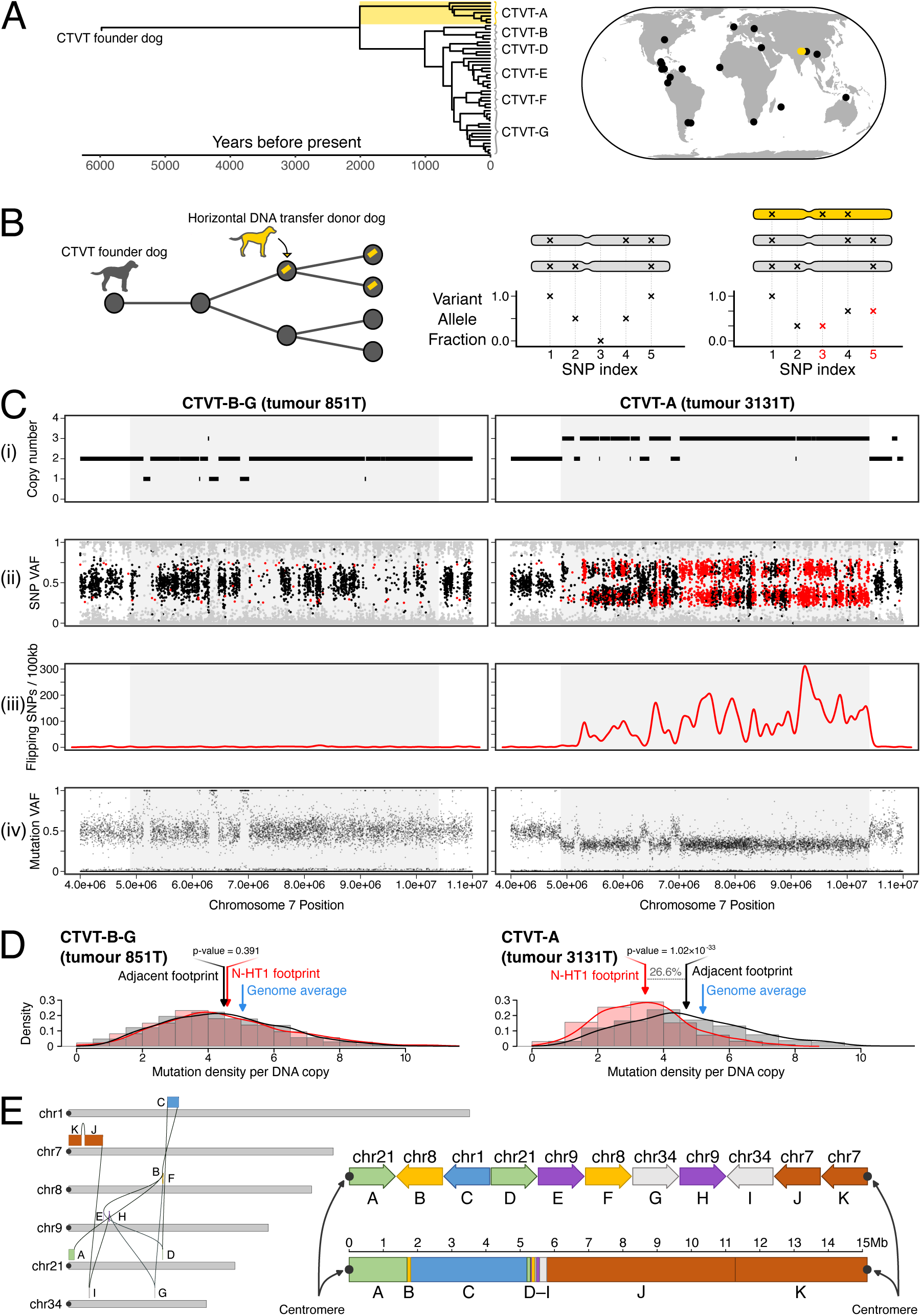
“Nuclear horizontal transfer 1” (N-HT1) is a horizontally transferred nuclear DNA element in CTVT-A. **(A)** Time-scaled phylogenetic tree and map illustrating evolutionary relationship and sampling locations, respectively, of 47 CTVT tumours analysed in this study. Gold identifies CTVT-A. Details and metadata available in Supplementary Figure 1 and Supplementary Table 1. **(B)** Schematic diagram depicting a screen for horizontal transfer of nuclear DNA in transmissible cancers. Left: CTVT (represented by filled circles) carries the constitutive genome of its original “founder dog” (dark grey), and is transmitted through transient hosts, generating a transmission tree (linked nodes). One such transient host (gold) acts as a donor (arrow) of genetic material (gold bar) which persists within child lineages in the tree. Right: focusing on genomic regions retaining both parental chromosome homologues (grey chromosomes above variant allele fraction plots), the screen searched for runs of “flipping Single-Nucleotide Polymorphisms (flipping SNPs)” (red) which flip from homozygosity to heterozygosity in a subset of tumours of a single cancer through the introduction of horizontally transferred DNA (gold chromosome). **(C)** Data corresponding to a 5.5 Mb interval of DNA horizontal transfer on chromosome 7 (N-HT1 segment J) for representative tumours belonging to CTVT-B–G (left, CTVT-G tumour 851T, horizontal DNA transfer absent) and CTVT-A (right, tumour 3131T, horizontal DNA transfer present). The horizontal transfer locus is in the centre of each panel marked with a grey background. Panels show (i) copy number, (ii) SNP variant allele fraction (VAF) corrected for tumour purity, (iii) density of SNPs flipping from homozygous to heterozygous state, and (iv) somatic mutation VAF corrected for tumour purity. In (ii), SNPs with heterozygous genotype in the majority of CTVTs are coloured black; those with homozygous genotype in the majority of CTVTs but heterozygous genotype in the sample shown (“flipping SNPs”) are coloured red; all other SNPs are coloured grey. Equivalent data for other implicated segments are available in Supplementary Figure 2. Plots in panels (ii) and (iv) have been limited to at most 20,000 individual points selected uniformly at random. **(D)** Distribution of somatic mutation density within a 5.5 Mb interval of DNA horizontal transfer (corresponding to the grey background shaded interval shown in (C)) in representative tumours belonging to CTVT-B–G (left, CTVT-G tumour 851T, horizontal DNA transfer absent) and CTVT-A (right, tumour 3131T, horizontal DNA transfer present). Red bars (“N-HT1 footprint”) represent distribution of somatic mutation density per 10 kilobases (kb) per chromosome copy within the shaded interval of chromosome 7 shown in (C) with mean indicated with red arrows. Grey bars (“adjacent footprint”) represents the same quantity within the region immediately adjacent to the shaded interval, with mean indicated with black arrows. P-values were obtained from two-sided T-tests for differences in means. Blue arrows (“genome average”) indicate copy number-normalised mean mutation density across all 10 kb windows in the genome for the tumour sample shown. **(E)** N-HT1 composition and structure. Left, a connection map of N-HT1 as inferred by structural variant analysis and long read DNA sequencing. N-HT1 comprises 11 blocks (A–K) derived from six chromosomes, running from the centromeric tip of chromosome 21 to the centromeric tip of chromosome 7. Canine chromosomes are represented by grey bars, and centromeres are represented by filled black circles. Right, relative orientation and sizes of blocks. The upper and lower representations are equivalent; the upper representation shows the block organisation schematically, with block orientation indicated with arrow direction, while the lower half shows the blocks to scale. Centromeres are represented by filled black circles.

Initial screens for host-to-tumour DNA exchange in transmissible cancers detected numerous instances of mitochondrial genome (mtDNA) horizontal transfer^16,22–25^. Indeed, the repeated observation of mtDNA capture in transmissible cancers has lent credibility to the idea that exchange of mtDNA between somatic cells is not an uncommon occurrence in multicellular organisms^26^. The detection of nuclear DNA horizontal transfer in transmissible cancers is not straightforward, however, and no clear evidence of this has yet been obtained. Here, we undertook a screen of 174 deeply sequenced tumour genomes belonging to the three known mammalian transmissible cancers to test the hypothesis that host-to-tumour horizontal transfer of nuclear DNA has contributed to their evolution.

### A screen for horizontal transfer of nuclear DNA

We analysed 47 CTVT tumour genomes sampled in 19 countries and selected to maximise representation of this lineage’s genetic diversity (Figure 1A, Supplementary Figure 1)^17,27,28^. Likewise, 78 DFT1 and 41 DFT2 tumour genomes were chosen to represent these cancers’ spatiotemporal ranges^29^. Tumour genomes were sequenced with short reads to a median depth of 77x alongside genomes of each tumour’s matched host (Supplementary Table 1).

We devised a screening approach to search for host-to-tumour nuclear DNA horizontal transfer (Figure 1B). Each transmissible cancer – DFT1, DFT2 and CTVT – arose from the cells of its respective founder animal. Although these three animals are themselves long-dead, their constitutive germline genotypes are clonally maintained in their transmissible cancers. Our intention was to screen tumours for deviations from clonality through the detection of germline genetic polymorphism.

Briefly, for each transmissible cancer (DFT1, DFT2, CTVT), we first selected genomic segments for which both of the respective founder animal’s parental chromosomal homologues were retained. This was evidenced by the presence of heterozygosity at germline single-nucleotide polymorphism (SNP) positions. Because we cannot access normal DNA from the three transmissible cancers’ founder animals, horizontal gene transfer encompassing regions which did not fulfil this criterion would be indistinguishable from simple loss-of-heterozygosity. Focusing on these selected segments, we screened for “flipping SNPs”: germline SNP loci whose genotypes were homozygous in some tumours, but heterozygous in other tumours belonging to the same transmissible cancer (DFT1, DFT2 or CTVT). Such shifts in genotype could individually be explained by mutation or localised homologous recombination. Extended runs of flipping SNPs, however, could signify the introduction of a non-parental DNA haplotype through a process of horizontal gene transfer (Figure 1B). The screen was designed to identify horizontal transfer events which occurred subsequent to the most recent common ancestor of each cancer’s analysed tumours, and which encompassed loci represented within the dog or Tasmanian devil reference genomes. Horizontal transfer events introducing additional copies of a locus (Figure 1B), as well as those replacing an existing copy, would theoretically be detectable under this approach.

The screen produced no evidence of DNA horizontal transfer in either of the Tasmanian devil transmissible cancers. In CTVT, however, we found regions of high flipping SNP density on chromosomes 1, 7 and 8 (Figure 1C, Supplementary Figure 2). The signal, whose length varied from 0.1 to 10 megabases (Mb), was confined to seven tumours belonging to a distinctive CTVT sublineage named “CTVT-A”. CTVT-A, which has so far only been detected in Nepal and India^17^, diverged about 2,000 years ago from its globally distributed sister sublineage, CTVT-B–G (Figure 1A, Supplementary Figure 1). Closer inspection of the genomic regions enriched for flipping SNPs revealed that the copy number in CTVT-A was, at all three loci, one integer step higher than that of CTVT-B–G tumours (Figure 1C, Supplementary Figure 2). These findings are consistent with the introduction of additional DNA haplotypes, encompassing three discrete genomic regions, to CTVT-A subsequent to its divergence from CTVT-B–G.

CTVT DNA accumulates somatic mutations with the passage of time^29^. It follows that CTVT constitutive DNA, present since the lineage’s origin, would be expected to have accrued more mutations than DNA captured by CTVT cells in a more recent process of horizontal transfer. Consistent with this, the density of mutations per DNA copy in CTVT-A tumours at the candidate horizontal transfer loci was on average 27 percent lower than that of flanking regions (Figure 1D, Supplementary Figure 3). Mutation density per DNA copy at these loci in CTVT-B–G tumours was, on the other hand, indistinguishable from flanking regions and similar to the genome average (Figure 1D, Supplementary Figure 3). Also of note, somatic mutations in the candidate horizontal transfer loci were invariably observed with variant allele fraction of 1/3 in regions of copy number 3 (Figure 1C, Supplementary Figure 2). This pattern is inconsistent with recent duplication of a parental chromosome, in which case mutations on 2/3 chromosome copies would also be observed, and is better explained by introduction of an additional horizontally transferred haplotype lacking somatic mutations.

We next explored whether the three detected candidate horizontal transfer loci on chromosomes 1, 7 and 8 were physically connected, thus potentially representing a single host-to-tumour DNA capture event in CTVT-A. Starting with the candidate locus on chromosome 7, one end of which terminated at this chromosome’s telocentric centromere, we detected a series of simple rearrangements which traversed the genome (Figure 1E, Supplementary Table 2). These connected not only the detected candidate loci, but also implicated additional segments on chromosomes 9, 21 and 34, ending at the centromere of chromosome 21 (Figure 1E). This configuration was supported by long read DNA sequencing (Supplementary Table 2). Inspection of the additional segments confirmed the presence of flipping SNPs specific to CTVT-A in most cases, although at a density insufficient for detection during the initial screen (Supplementary Figure 2). The additional loci also presented other hallmarks of horizontal gene transfer, including integer step changes to higher copy number in CTVT-A relative to CTVT-B–G, absence of somatic mutations with 2/3 variant allele fraction in regions of copy number 3, and reduced mutation density per DNA copy specific to CTVT-A (Supplementary Figure 3). The magnitude of this reduction was similar across implicated loci (Supplementary Figure 3). These findings uncover a 15-Mb DNA element, pieced together from 11 assorted fragments of six chromosomes, and predicted to be flanked on both ends by centromeric sequence (Figure 1E, Supplementary Table 3). This fragment, which we have termed “nuclear horizontal transfer 1”, N-HT1, was taken up and retained by CTVT-A cells through a process of DNA horizontal transfer.

### Cytogenetic organisation of the horizontally transferred DNA element

Despite its di-centromeric structure, N-HT1 is stably present in a single copy in tumours belonging to CTVT-A. We investigated the physical organisation of N-HT1 within CTVT-A nuclei using metaphase fluorescence *in situ* hybridisation. Three bacterial artificial chromosome probes were selected from genomic regions spanned by N-HT1, including a probe from a locus homozygously deleted on CTVT parental chromosomes and expected to be uniquely represented on N-HT1 (Figure 2A). These were hybridised to CTVT-A metaphases as well as to control metaphases from CTVT-B-G tumours. The three probes co-localised in CTVT-A metaphases, but not in CTVT-B-G metaphases, revealing that N-HT1 forms the short arm of a small submetacentric chromosome (Figure 2B, Figure 2C, Figure 2D). This implies that N-HT1 has been incorporated onto a CTVT chromosome through a centromeric fusion. Its second centromere has presumably been inactivated^30^, producing the stable structure that we observe.

**Figure 2:**
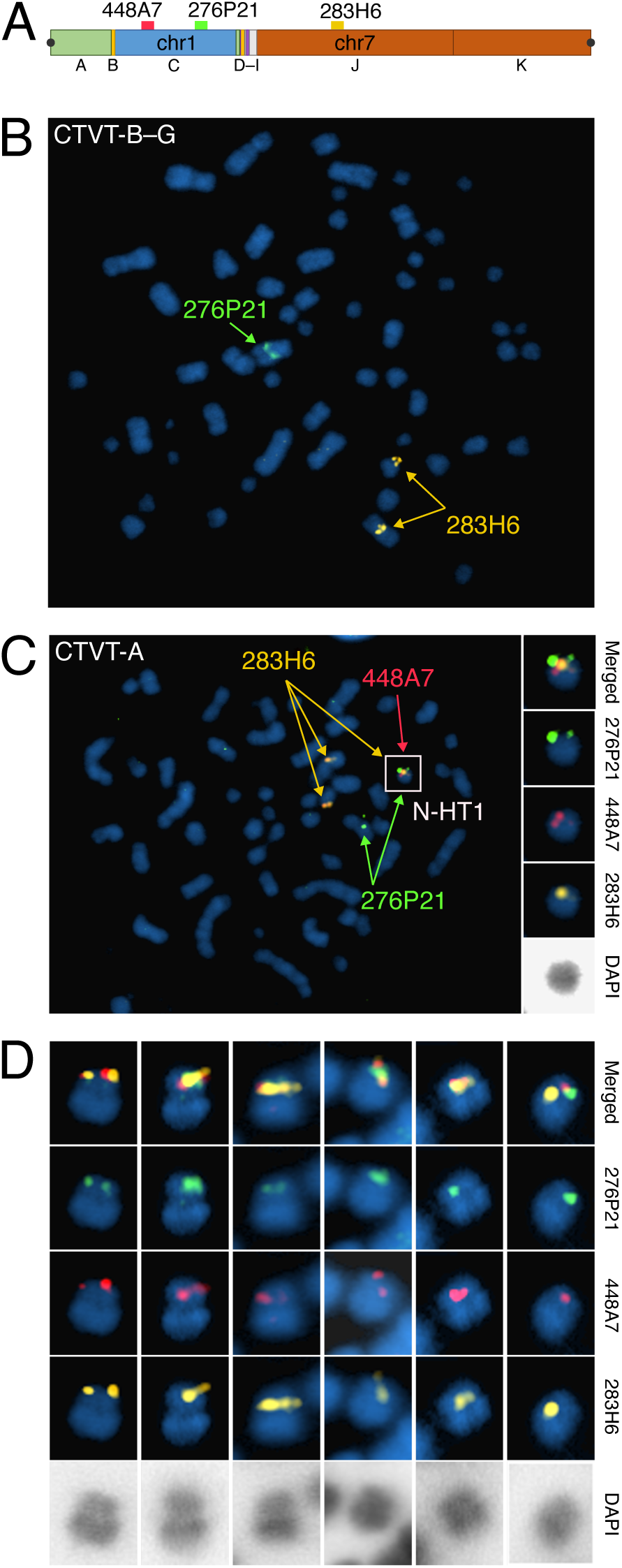
Cytogenetic visualisation of N-HT1. **(A)** Schematic representation of N-HT1 with mapping locations of bacterial artificial chromosome probes CH82-448A7 (red), CH82-276P21 (green) and CH82-283H6 (gold) shown. The locus homologous to probe CH82-448A7 (red) is homozygously deleted on CTVT parental chromosomes, and N-HT1 is expected to be its only mapping location in CTVT. Centromeres are represented by black circles. **(B)** Fluorescently labelled probes hybridised to metaphase chromosomes of a CTVT-B-G tumour, 3838T. CH82-283H6 (gold) maps to two chromosomal loci, CH82-276P21 (green) to a single chromosomal locus, and CH82-448A7 (red) is absent. 100x magnification. **(C)** Fluorescently labelled probes hybridised to metaphase chromosomes of a CTVT-A tumour, 3751T. In addition to the CTVT parental chromosome mapping locations identified in (B), the three probes co-localise to a single chromosome that is identified to contain N-HT1 (white box). 3x enlargements of the chromosome containing N-HT1 with merged probe channels, individual probe channels and inverted DAPI are shown on right. 100x magnification. **(D)** Six additional examples of the N-HT1 chromosome hybridised with the three probes illustrating that N-HT1 forms the short arm of a small submetacentric chromosome. Images show merged probe channels, individual probe channels and inverted DAPI. 3x enlargements from 100x magnification. Images are from tumour 3751T (from left to right, images 1, 2, 3, 5 and 6) and tumour 3770T (from left to right, image 4).

### Origin of horizontal DNA transfer in CTVT

The specificity of N-HT1 to tumours belonging to CTVT-A suggests that it was captured by this sublineage after CTVT-A diverged from CTVT-B–G approximately 2,000 years ago^30^. We used somatic mutation density to estimate the time since N-HT1’s incorporation into CTVT. Focusing on cytosine-to-thymine (C>T) substitution mutations occurring at CpG dinucleotide sites, which accumulate at a constant rate^31^, we estimated a density of 2.21×10^−3^ (95% CI: 2.16–2.25×10^−3^) mutations per site per CTVT parental chromosome copy in CTVT-B–G tumours at the genomic regions spanned by N-HT1. Assuming that CTVT-A parental chromosomes carry a similar mutation density to those of CTVT-B–G, the mutation density attributable to N-HT1 was estimated as 6.89×10^−4^ (range 5.82–7.96×10^−4^) mutations per site per chromosome copy (Figure 3A). Assuming a CTVT mutation rate of 6.87×10^−7^ mutations/site/year per diploid genome^17^ (3.435×10^−7^ mutations/site/year per chromosome copy), this mutation density corresponds to 2,006 (95% CI: 1,695–2,317) years ago. The most parsimonious interpretation of these findings is that N-HT1 was acquired by CTVT-A shortly after it split from CTVT-B–G (Figure 3A).

**Figure 3:**
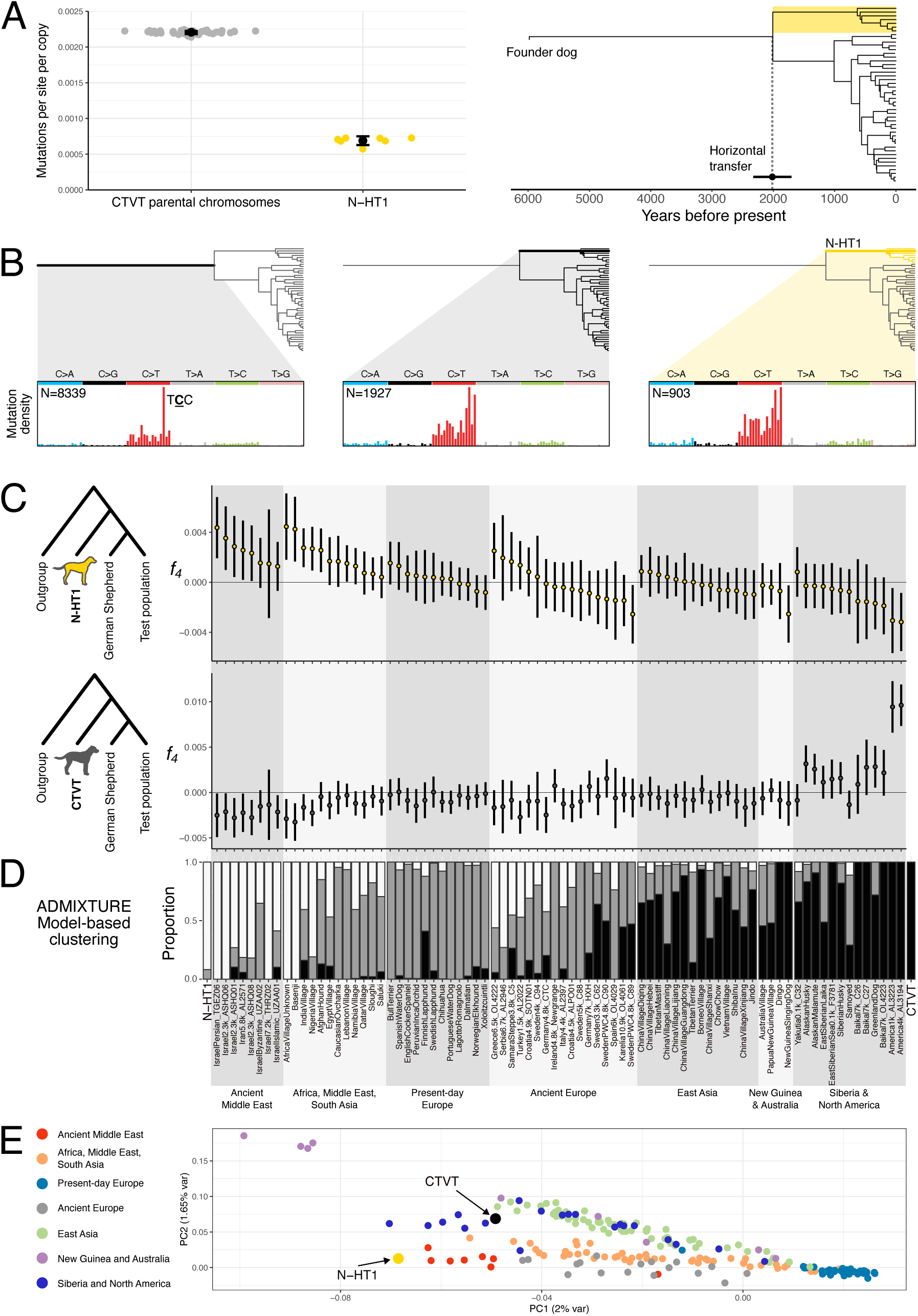
Timing of horizontal transfer and origin of N-HT1. **(A)** Estimation of the relative timing of N-HT1 horizontal transfer. Left, cytosine-to-thymine substitution mutations per CpG dinucleotide site per DNA copy for CTVT parental chromosomes and N-HT1. For “CTVT parental chromosomes”, each point represents mutation density per site per copy within the intervals corresponding to the N-HT1 footprint for an individual tumour. Points for “N-HT1” represent mutation density in CTVT-A tumours after subtracting mutation counts attributable to CTVT parental chromosomes (estimated using mutation counts observed in CTVT-B–G) from observed mutation counts. Bars show the mean and its 95% confidence interval for each group. Right, N-HT1 acquisition time estimated from mutation densities. Estimate is plotted as a black point superimposed on the CTVT time-calibrated phylogeny. The horizontal black line shows 95% confidence intervals around the point estimate calculated by propagating the standard errors in the N-HT1 group. CTVT-A is highlighted with a gold box. **(B)** Mutation spectra for specific lineages in the CTVT phylogenetic tree. Mutations shared by CTVT-A and CTVT-B–G but absent from N-HT1 (left) have a large proportion of T<u>C</u>C to T<u>T</u>C mutations (mutated base underlined). Mutations unique to CTVT-A and absent from N-HT1 (centre), and unique to N-HT1 (right) show similar mutation spectra. Mutation density plots show proportion of mutations at each of 96 trinucleotide sequence contexts, coloured by mutation type; fully labelled plots are presented in Supplementary Figure 4. N, number of mutations within each subset. **(C)** *f_4_*-statistics calculated for quartets of canid populations in the form *f_4_*(Outgroup, N-HT1; German Shepherd Dog, Test population) (upper row) or *f_4_*(Outgroup, CTVT; German Shepherd Dog, Test population) (lower row) from germline transversion SNPs occurring in genomic intervals spanned by N-HT1. Andean fox was used as outgroup and Test populations are listed on *x*-axis (*x*-axis labels are shared with (D)). Trees represent the null hypothesis of the F_4_ test: that the groups (Outgroup, N-HT1) and (German Shepherd Dog, Test population) (upper panel) or the groups (Outgroup, CTVT) and (German Shepherd Dog, Test population) (lower panel) form clades with respect to each other. *f_4_*-statistics significantly different from zero reject this hypothesis; positive values of *f_4_* suggest that (N-HT1, Test population (upper panel)) or (CTVT, Test population (lower panel)) are sufficiently closely related that they form a clade with respect to (Outgroup, German Shepherd Dog). Error bars represent *f_4_* ± 3 standard deviations, estimated by block 1jackknife sampling. **(D)** Model-based clustering (ADMIXTURE) for the same population groups as shown in (C). Data represent the admixture proportions into the labelled populations from three underlying latent ancestry streams denoted with white, grey and black. Values are pooled in cases where the labelled population is represented by multiple samples. **(E)** Principal component analysis performed on genotypes of 124,209 transversion single-nucleotide polymorphisms occurring within region spanned by N-HT1. Analysis was performed on genotypes from 759 dogs, N-HT1 and CTVT. Only samples belonging to the populations described in (C) and (D) are shown. Data are plotted according to their projection onto first and second principal components, and the proportion of variance explained by these is indicated. Details of dog populations used in (C), (D) and (E) are summarised in Supplementary Table 8.

Mutation signatures provide additional evidence that N-HT1 was acquired relatively recently in CTVT’s history. Mutations acquired before, but not after, the most recent common ancestor of sampled tumours are highly enriched for a distinctive pattern known as “signature A” characterised by C>T mutations at a T<u>C</u>C context (Figure 3B, mutated base underlined, Supplementary Figure 4)^31,32^. Mutations phased to N-HT1 do not show signature A, implying that N-HT1 was captured after exposure to signature A ceased (Figure 3B).

N-HT1 is a rearranged genomic fragment belonging to a dog that lived around the start of the Common Era. We know that this dog became infected with CTVT, developing its characteristic genital tumours. It subsequently passed its CTVT cells, now modified to include a small fragment of its own DNA, to its mating partners. The DNA fragment which this dog donated, however, bears further testimony to this animal’s identity in its genetic sequence. We inferred the N-HT1 genotype at 124,209 polymorphic SNP positions by subtracting counts of CTVT parental alleles, themselves determined using CTVT-B–G tumour sequences, from observed CTVT-A allele frequencies. Next we employed *f_4_*-statistics to explore genetic relationships among N-HT1 and CTVT genotypes and those of 80 modern and ancient dog populations^32,33^ (Figure 3C). As expected, CTVT shared excess derived alleles with precontact North American dogs, the population from which CTVT itself originated^32^. N-HT1 was distinct from CTVT, however, and shared genetic drift with ancient Middle Eastern dogs as well as modern dogs sampled in Africa, the Middle East and South Asia^32,33^. Model-based clustering^22,35^ and principal component analysis further supported this finding (Figure 3D and Figure 3E).

Dogs have a complex history characterised by migration and several major admixture episodes^32,33^. Although we can access only a 15 Mb fragment of the N-HT1 donor dog’s genome, it suggests that this animal belonged to a dog population that occurred in the Middle East, and possibly further afield across the wider South and Central Asian region, Africa and Europe, around the start of the Common Era. This population shows genetic continuity with dogs which today inhabit the Middle East, Africa and South Asia^32^. Given that CTVT-A itself has been detected only in Nepal and India, CTVT-A may not have strayed far from its present location since it diverged from CTVT-B–G and acquired N-HT1 some 2,000 years ago.

Of note, CTVT-A carries a unique mtDNA haplotype, known as mtDNA-HT4, that was captured through a horizontal transfer event 1,723–2,362 years ago^36^. It is thus plausible that N-HT1 and mtDNA-HT4 were acquired by CTVT-A in a single transfer from the same donor dog. Phylogenetic analysis of canine and CTVT mtDNA does not, however, indicate a genetic relationship between mtDNA-HT4 and mtDNA of ancient Middle Eastern dogs^32^ (Supplementary Figure 5). Indeed, the latter themselves carry a number of unrelated mtDNA haplotypes^32^. This analysis confirms that several mtDNA haplotypes were segregating in the dog population to which N-HT1 belonged and does not discount the possibility that an individual donor dog contributed both nuclear and mitochondrial DNA to CTVT-A in a single horizontal transfer event.

### Gene expression from the horizontally transferred DNA element

N-HT1 encodes 133 intact protein-coding genes as well as 10 gene fragments truncated by segment boundaries (Supplementary Table 4). We investigated expression of these by screening for informative variants that could distinguish N-HT1 transcripts from those of CTVT constitutive chromosomes as well as those of each tumour’s individual matched host (Figure 4A). Using RNA sequencing, we quantified relative expression of N-HT1-encoded transcripts for the 73 genes possessing such informative alleles (Supplementary Table 5). These data revealed that N-HT1 is transcriptionally active, and that its pattern of gene expression resembles that of CTVT constitutive chromosomes rather than that of tumour-infiltrating host cells (Figure 4B, Supplementary Table 5). Thus, exposure to the CTVT cellular environment was sufficient to reprogramme this DNA fragment’s transcriptional repertoire.

**Figure 4:**
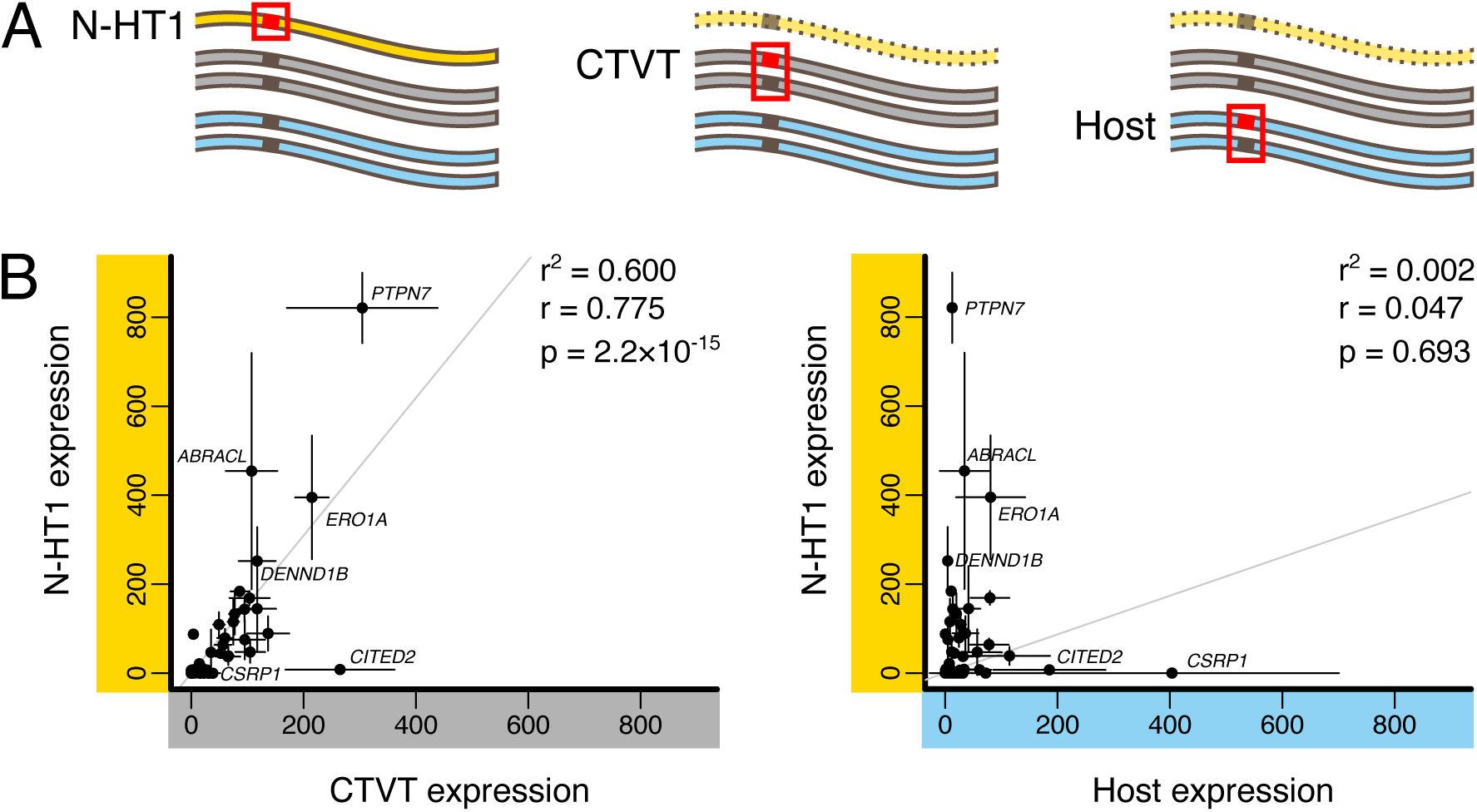
Expression of genes encoded on N-HT1. **(A)** Allelic deconvolution of bulk tumour RNA sequencing data was used to quantify transcript expression for genes encoded on N-HT1. Upper diagrams depict DNA carrying alleles that are informative about N-HT1 (left), CTVT parental chromosomes (centre) and matched host (right). Informative alleles are represented by red squares and boxes. N-HT1 is made partially transparent to indicate that N-HT1 may or may not be present in genotype combinations informative on CTVT and host. **(B)** Transcript read counts of informative alleles from normalised bulk tumour RNAseq were used to quantify N-HT1, CTVT and matched host gene expression (from tumour-infiltrating host cells) in tumours carrying the illustrated informative genotype combinations. Scatterplots show the relationship between the mean allelic expression per chromosome copy for N-HT1 and CTVT (left), and for N-HT1 and tumour-infiltrating matched host (right). Each point represents the mean relative expression per chromosome copy, estimated using normalised transcript read counts, for each of 73 genes for which informative expression data were available. Gene names are indicated for a subset of genes. Vertical and horizontal bars around each point show 95% confidence intervals around these means. Least squares regression trend lines are shown, together with Pearson’s correlation coefficient (r) and r^2^ values, and associated p-value as computed by the R function ‘cor.test’. Source data are available in Supplementary Table 5.

None of the N-HT1-encoded genes possess known relevant dosage-dependent oncogenic roles in cancer. It is worth noting, however, that one gene, *ARFGEF3*, was rescued from a null state by the introduction of N-HT1. *ARFGEF3*, which encodes a guanine nucleotide exchange factor, was biallelically inactivated in the common ancestor of CTVT-A and CTVT-B–G through a combination of nonsense mutation and deletion (Supplementary Table 6). N-HT1 complemented this loss, although the N-HT1 copy has apparently also subsequently undergone inactivation by acquisition of an independent nonsense mutation (Supplementary Table 6). Similarly, 189 kilobases of DNA homozygously deleted in the CTVT-A and CTVT-B–G common ancestor was reinstated in CTVT-A by N-HT1, although the affected loci do not encode protein-coding genes (Supplementary Table 7). Overall, these data provide no evidence that the introduction of N-HT1 significantly altered CTVT cellular function in the long term.

## Discussion

We can only speculate as to the series of events which created N-HT1 and conveyed it into CTVT. N-HT1 is highly internally rearranged, suggesting that it was formed by haphazard repair of fragmented DNA prior to its introduction to CTVT. The repeated representation of segments of only six chromosomes within N-HT1 hints at the possibility that these were physically segregated within the donor cell, perhaps inside a micronucleus. The dicentric organisation of N-HT1 implies that this was an unstable linear or circular element within the donor cell. Indeed, the aberrant internal structure of N-HT1 suggests that this DNA fragment occurred within a cell that was not healthy. Perhaps this cell was dying or even dead, with the possibility that N-HT1 was sequestered within an apoptotic body^36–39,38–41^.

Regardless of the state of the donor cell, N-HT1 came to enter a CTVT cell. This may have involved cell fusion followed by expulsion of the remainder of the donor cell genome. Alternatively, donor cell material may have been taken up by phagocytosis, with N-HT1 subsequently escaping lysosomal degradation and double-stranded DNA sensing innate immunity^42^ and entering the nucleus. Other transfer mechanisms are also possible. It seems plausible that a circular structure, which would have lacked the double-stranded DNA breakpoint ends conferring susceptibility to nuclease degradation and DNA damage checkpoint activation in the recipient cell^43^, may have stabilised N-HT1 during this process. Once within the CTVT nucleus, a centromeric fusion event incorporated N-HT1 onto an existing CTVT chromosome to produce a small novel chromosome. N-HT1 subsequently persisted for centuries, passing through innumerable canine hosts, and survives to this day within the tumours of dogs roaming the streets of Nepal and India.

This work suggests that host-to-tumour horizontal transfer of nuclear DNA is rare in dog and Tasmanian devil transmissible cancers, although it is important to note that the screen was limited by availability of genetic markers and that it surveyed only the lines of cells that both contributed to tumour transmission and which occurred subsequent to the most recent common ancestors of each cancer’s sampled tumours. The latter implies that any DNA exchange that contributed to tumour initiation would not have been identified. Nevertheless, our survey of centuries of CTVT evolution, as well as decades of Tasmanian devil DFT1 and DFT2 evolution, found just one instance of host-to-tumour nuclear DNA horizontal transfer. Moreover, it appears unlikely that this single detected event was adaptive for CTVT. N-HT1 does not encode genes with obvious relevant links to oncogenesis. Furthermore, CTVT-A, which hosts N-HT1, is rare and spatially restricted, in contrast to its globally distributed CTVT-B–G sister sublineage. We conclude that host-to-tumour horizontal transfer of nuclear DNA has not been a relevant driving force contributing to mammalian transmissible cancer evolution, and that the single event that we detected is a selectively neutral “passenger”.

Despite this, we emphasise that the incidence and importance of nuclear horizontal DNA transfer may vary among cancers. Although long recognised as a potential source of genetic variation in cancer^44^, technical difficulties have precluded large-scale screening for its contribution to human tumours. Detection of host-to-tumour horizontal gene transfer has thus far been limited to experimental models^45^ and suggestive observations in cancers occurring in transplant recipients^46^. The increased genetic resolution afforded by advances in single cell sequencing and long-read sequencing now presents opportunities for examining the frequency of this mode of DNA variation across large numbers of cancer genomes. The potential of horizontal gene transfer to drive mutation reversion, for instance in response to therapeutic pressure^44^, is particularly worth examining; although apparently not of selective consequence, the complementation of biallelically inactivated *ARFGEF3* by N-HT1 illustrates this concept. Of note, it is not unlikely that host-to-tumour nuclear DNA horizontal transfer may in some cases have negative fitness consequences for the cancer cell, perhaps contributing to the rarity of this phenomenon. However, as with other modes of mutation in cancer, most instances of horizontal gene transfer are likely to be selectively neutral passengers^28,47^, and our study has uncovered one such event which has immortalised a piece of DNA from a long-dead dog. Identifying those cases where horizontally transferred DNA drives positive selection may reveal new conceptual and therapeutic avenues in cancer biology.

## Supplementary information and data accession

This paper includes Supplementary Figures 1-5 and Supplementary Tables 1-8. Supplementary Data 1-5 are available on Zenodo (https://doi.org/10.5281/zenodo.7214808). Sequence data are available on the European Nucleotide Archive (accession in progress).

## Acknowledgements

We thank Andrew Chan, George Carnell, Paul Edwards, Michelle Smith and Robin Moll for technical assistance. We are grateful to Debbie Sabin and the Histology team at the Department of Veterinary Medicine, University of Cambridge. We thank Alexander Sampson for helpful comments on the manuscript. We acknowledge sequencing, IT and administrative staff at the Wellcome Sanger Institute for their support. We thank the following for their help obtaining samples for this project: Rakesh Chand, Luca Schiavo, Eric Davis, students from St. George’s University (True Blue, Grenada, West Indies) who assisted with sample collection, staff and volunteers at World Vets (Gig Harbor, USA), and staff at Animal Nepal (Kathmandu, Nepal). This work was supported by Wellcome (102942/Z/13/A, 222551/Z/21/Z), BBSRC (BB/Y514299/1), and grants from King’s College, Cambridge, and St John’s College, Cambridge.

## Author Contribution

E.P.M. conceived the project. K.G. performed the analysis with input from E.P.M., A.B.-O and M.R.Stammnitz. The manuscript was written by E.P.M. and K.G. with contributions from A.B.-O., A.S., M.R.Stammnitz, J.W. and M.R.Stratton. J.W., A.S., J.C., S.B. and M.H. processed the samples, K.H., D.M., J.B. and E.D. performed laboratory investigation. K.M.A., M.V.B., A.M.C., K.F.d.C., E.M.D., I.A.F., A.H., N.I., R.K., D.K., M.L.-P., A.M.L.Q., M.M., W.N., F.P.-O., Y.P., K.P., J.C.R.-A., J.F.R., S.K.S., S.S., L.J.T.M., M.T., Sa.T., Su.T., M.G.v.d.W., and A.S.W.-M. collected clinical samples. All authors reviewed and approved the manuscript.

## Materials and methods

### Sample preparation and sequencing

#### Sample collection and nucleic acid extraction

Dog sample collection was approved by the Department of Veterinary Medicine, University of Cambridge, Ethics and Welfare Committee (reference number CR174). Tumour and host tissue was collected into RNAlater (Invitrogen, Carlsbad, CA, USA), with the exception of sample 2T, which was stored in ethanol. Genomic DNA for short read sequencing was extracted using the Qiagen DNeasy Blood and Tissue extraction kit (Qiagen, Hilden, Germany), and total RNA was extracted using the Qiagen AllPrep Universal Kit (Qiagen, Hilden, Germany). High molecular weight DNA was extracted from two CTVT tumours using the Qiagen MagAttract High Molecular Weight DNA Kit (Qiagen, Hilden, Germany). Dog sample metadata are available in Supplementary Table 1. A publicly available Tasmanian devil data set was used in this study^29^.

#### Whole genome sequencing and alignment

Standard whole genome sequencing libraries with insert sizes ranging from 450 to 530 base pairs (bp) were prepared from genomic DNA extracted from 47 CTVT tumours, as well as from 46 CTVT matched host dogs (Supplementary Table 1). Whole genome sequencing with 125 bp or 150 bp paired end reads was performed using Illumina HiSeq 2000, HiSeq X Ten or NovaSeq S4 instruments (Illumina, San Diego, CA, USA) (Supplementary Table 1). Adaptor sequences were trimmed using biobambam^44^ (versions 2.0.17-2.0.79) and reads were aligned to CanFam3.1^45^ (ENA accession: GCA_000002285.2) using BWA-mem^48^ versions ranging between 0.7.12-r1039 and 0.7.17-r1188, using settings `-p -Y -K 100000000 -T 30` (HiSeq 2000 samples only: `-K 120000000 -T 0`). PCR duplicates were marked using biobambam^49^ (versions 2.0.17-2.0.79). Publicly available canine whole genome sequencing data^44^ were processed and aligned in parallel with the new data. Dog sample metadata are available in Supplementary Table 1. Tasmanian devil DNA was sequenced and aligned as described^29^.

#### RNA sequencing and alignment

RNA extracted from CTVT tumours was used to prepare random primed ribosomal RNA depleted RNAseq libraries with insert size 240-280bp as described^50,51^. Approximately 314 million 100 bp paired end reads were sequenced from each library. Adaptors were trimmed using biobambam^52^ 2.0.79, and reads were aligned to CanFam3.1^51^ (ENA accession: GCA_000002285.2) using STAR^53^ version 2.5.2b, with settings

`--runMode alignReads \`

`--runThreadN 12 \`

`--genomeLoad NoSharedMemory \`

`--outStd BAM_Unsorted \`

`--outSAMtype BAM Unsorted \`

`--outSAMstrandField intron Motif \`

`--outSAMattributes NH HI NM MD AS XS \`

`--outSAMunmapped Within KeepPairs \`

`--outFilterIntronMotifs RemoveNoncanonicalUnannotated \`

`--chimSegmentMin 0 \`

`--chimJunctionOverhangMin 20 \`

`--chimOutType WithinBAM \`

`--sjdbOverhang 74 \`

`--quantMode GeneCounts`

using the Ensembl^54,55^ 95 CanFam3.1 transcriptome as a splice junction DB (`--sjdbGTFfile`). Alignment files were sorted and PCR duplicates were marked using biobambam^56,57^ 2.0.79.

### Long read DNA sequencing and alignment

High molecular weight DNA from two CTVT tumours was sequenced using the Pacific Biosciences HiFi Revio sequencing platform (Pacific Biosciences, Menlo Park, United States). HiFi reads were aligned to CanFam3.1^58^ (ENA accession: GCA_000002285.2) using pbmm2 v1.13.1^50,51^ with 8 aligning threads and 4 sorting threads. Sample metadata are available in Supplementary Table 1.

### Single base substitution and indel variant calling and annotation

Single base substitution and indel variants were called using Platypus v0.8.1^57^ as part of the Somatypus v1.3 variant calling pipeline, as previously described^59^. Tasmanian devil variants were further filtered as described in Stammnitz *et al.* (2023).

Dog variants were further filtered as follows:

- *Unplaced contig filter:* Variants not located on chromosomes 1–38 or X were removed.
- *Host coverage filter:* Variants with a median coverage among hosts of fewer than 10 reads or more than 200 reads were removed.
- *Low read support filter:* Variants that occur with 1 to 4 supporting reads in at least one host, and never occur with 5 or more reads in any one sample (tumour or host) were removed, as they are likely to represent low-frequency sequencing or alignment errors.

Next, dog variants were defined as either “somatic” or “germline”:

- Somatic: Variants are categorised as somatic (tumour-only) if they occur with three or more supporting reads in one or more tumours and have a variant allele fraction (VAF) less than 0.25 in all hosts.
- Germline: Variants are categorised as germline (host and tumour) if they have VAF above 0.25 in any host.

Variants that did not fulfil the definition of either somatic or germline were discarded.

Next, a preliminary phylogenetic tree was generated with somatic variants using IQ-TREE v2.0.7^60^, with model GTR+G{4}^61^. A “low tumour support phylogenetic filter” was applied as follows:

- *Low tumour support phylogenetic filter:* The intention was to discard variants with low support, while retaining low VAF variants in closely related tumours which may represent shared subclonal variants. First, somatic variants which were supported by 1 to 4 reads in each of at least 2 tumours, and never supported by 5 or more reads in any one tumour, were collected. These were retained only if the tumours which showed read support were phylogenetically related. This was defined as requiring only a single ancestral event to have occurred to create the observed presence pattern at the tips of the preliminary tree. Ancestral reconstruction was performed by maximum parsimony using Phangorn^62^.

Somatic variants were further filtered as follows:

- *Excessive host support filter:* Somatic variants with 5 or more reads in any single host were discarded as having excess host support.
- *Repetitive region filter:* Somatic variants within 5 bp of annotated repeat regions (simple, low-complexity or tandem repeats) were discarded.
- *Strand bias filter:* Somatic variants with 20% or less of total supporting reads on either strand were discarded.

Tumour purity was estimated as previously described^29^. Briefly, the mode of the distribution of VAF for somatic substitution mutations was determined, and purity was estimated as *p* = 2 × *mode*(*VAF*), where *p* is purity. Finally, the following filter was applied:

- *Unmatched tumour filter:* Somatic variants unique to a tumour with no matched host, and with VAF≤ 0.3 ×purity were discarded.

Variants were annotated using the Ensembl Variant Effect Predictor version 104^64^.

### CTVT group assignment

The CTVT phylogenetic tree from Baez-Ortega *et al.* (2019)^43^ was divided into seven nested monophyletic groups, designated CTVT-A through -G (Supplementary Figure 1). Tumours included in the current study were selected to provide representatives from each group, but CTVT-C was not included (group C corresponds to “Group 1 – India (Kolkata)” in Baez-Ortega *et al.* (2019)^45^).

### Copy number estimation

Copy number was estimated for each clone separately (CTVT, DFT1 and DFT2), using the method described in Stammnitz *et al.* (2023)^29^. Only scaffolds assigned to chromosomes were considered. CTVT ploidy was estimated using a grid search approach^29^; all tumours were close to diploid (Supplementary Table 1). Copy number data are available in Stammnitz *et al.* (2023)^29^ and in Supplementary Data 2.

### Purity correction of variant allele fraction

Variant allele fraction (VAF) purity correction was performed on germline single-nucleotide polymorphisms (SNPs) and somatic mutations in tumours for which matched host was available, by the method below.

An estimate of the probability that a read derives from the host genome, at a given position, is obtained as:

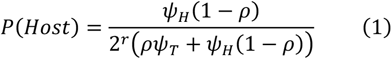

where *r* is the log read ratio (logR) at the position, *ρ* the sample purity, and *ψ_H_*, *ψ_T_* the sample ploidy for the host and tumour, respectively. The log read ratio, logR, is the base-2 logarithm of the normalised read coverage of the tumour divided by the normalised read coverage of the host. Methods for computing the logR are described in Stammnitz *et al.* (2023)^29^.

Assuming host ploidy of 2, this simplifies to:

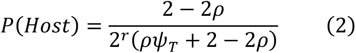

An estimate of the probability that a host read carries the alternative allele is directly obtained from the VAF of the position in the matched host:

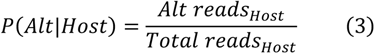

The total number of reads derived from the host (*K*) can be estimated as:

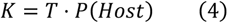

where *T* is the total number of reads observed in the tumour sample at this position.

Likewise, the total number of alternative-allele reads derived from the host (*L*) can be estimated as:

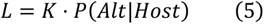

This implies that the total number of reads due to the tumour is *T* − *K*, and the total number of alternative-allele reads due to the tumour is *A* − *L*, where *A* is the number of alternative-allele reads observed in the tumour sample at this position, and hence an estimate of the VAF for an uncontaminated tumour is given by:

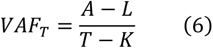

where *A* is the number of alternative-allele reads observed in the tumour sample at this position. In practice, the corrected tumour VAF is estimated by least-squares optimisation of

*K* and *L*, subject to non-negativity constraints on the quantities *L* (alternative-allele reads in host), *A* − *L* (alternative-allele reads in tumour), *K* − *L* (reference-allele reads in host) and *T* − *K* − *A* + *L* (reference-allele reads in tumour), incorporated as Lagrangian multipliers. The implementation of the optimisation is included in the accompanying R code (https://github.com/TransmissibleCancerGroup/NuclearHorizontalTransfer).

### Flipping SNP screen

Examining each tumour individually, each germline SNP was classified as “heterozygous”, “homozygous” or “unclassified”. Heterozygous SNPs were those whose purity-corrected VAF which fell within one of the intervals described below. Unclassified SNPs were those falling in segments of copy number 0 or 1. All other SNPs were classified as homozygous.

| Copy number | Interval lower bound | Interval upper bound |
| --- | --- | --- |
| 2 | 0.25 | 0.75 |
| 3 | 0.2 | 0.8 |
| 4 | 0.175 | 0.825 |
| 5 | 0.15 | 0.85 |
| 6 | 0.12 | 0.88 |
| $\geq 7$ | 0.1 | 0.9 |

Next, each SNP was given a “majority status”. For each SNP, let N_Het_ represent the number of tumours within the given lineage (CTVT, DFT1 or DFT2) with an assignment of “heterozygous”, and N_Hom_ represent the number of tumours within the given lineage (CTVT, DFT1 or DFT2) with an assignment of “homozygous”. If N_Het_>N_Hom_, then the SNP was assigned as “majority heterozygous”. If N_Hom_>N_Het_, the SNP was assigned as “majority homozygous”. Unclassified SNPs, with copy number 0 or 1, did not contribute to majority status.

Considering each tumour genome individually, a SNP whose genotype (heterozygous, homozygous) conflicted with the majority state for its lineage (CTVT, DFT1, DFT2) was defined as “flipping”.

Next, we calculated flipping SNP density. For each tumour, we considered only the subset of the genome occurring in copy number state 2 or higher, and we collated all SNP positions in this subset into a list. Flipping SNP density at each position in each tumour was calculated as the proportion of flipping SNPs within a window consisting of the 500 preceding and 500 following SNPs in the list (i.e. considering only those SNP positions occurring at copy number state 2 or higher in a given tumour), as well as the current SNP (1001 SNPs considered in total). Peaks in this density were automatically selected by calculating the Cook’s distance^67^ for each SNP, and selecting SNPs for which the Cook’s distance exceeded thirty times the mean Cook’s distance. These peaks were further filtered by requiring that the density should exceed the median flipping SNP density for that tumour plus five times the Median Absolute Deviation (MAD) of the density. Flipping SNP peaks outputted using this screen were then visually inspected for possible horizontal gene transfer. The flipping SNP screen initially identified regions of horizontal transfer on chromosomes 1, 7 and 8, corresponding to N-HT1 segments B, C, F, J and K. These segments were confirmed and the remaining ones added by structural variant detection, as described below.

### Structural variant detection

Structural variants were inferred using SvABA^68^, and the MSG pipeline^29^, which uses Manta as its structural variant caller^50^.

Starting with the five N-HT1 segments detected through the flipping SNP screen (B, C, F, J and K), we obtained structural variants that coincided with copy number change points flanking these segments. As expected, these were specific to CTVT-A. This was followed by a “walking” approach which linked N-HT1 segments with one another by following the inter-chromosomal rearrangements detected by Manta and SvABA.

Junction points between the eleven detected segments of N-HT1 were inferred to base-pair resolution by combining the output of Manta and SvABA with realignments made using svviz2^52^, and visual inspection using IGV^69^ (Supplementary Table 2A and Supplementary Table 3).

We validated the structure and breakpoint sequences of N-HT1 using PacBio HiFi long sequence reads. We identified 69 reads that spanned more than one junction point (Supplementary Table 2B).

N-HT1 segment A involves a genomic region at the centromeric tip of chromosome 21 that has undergone complex and polymorphic gains in copy number across the CTVT lineage. Part of this segment is present at copy number 4 in CTVT-A. We confirmed that the N-HT1 haplotype in this region is present in single copy by determining that somatic mutations with VAF 2/4 in this segment in CTVT-A are invariably shared with CTVT-B–G, indicating that the duplication involving this region in CTVT-A was acquired prior to the CTVT-A-CTVT-B–G most recent common ancestor, and therefore could not have involved N-HT1.

### Time-resolved phylogenetic inference

We used BEAST v1.10.4^70^ to construct a time-calibrated phylogeny for 47 CTVT tumours, following the procedure used in Baez-Ortega *et al.* (2019)^71^. Following the published procedure, the data used were C-to-T mutations occurring at CpG sites in the CanFam3.1 reference^45^, restricted to the exome (Ensembl gene build v104). This led to a matrix of 48 taxa (47 tumours plus idealised reference sequence), with 37,701 variable sites, with an additional 3,685,744 constant sites encoded in a `constantPatterns` block in the BEAST XML. The model and its priors were specified as follows: a fixed-population coalescent tree prior, with a “oneOnX” prior on population size; a Jukes-Cantor substitution model^49^ with gamma distributed rates (approximated by four discrete categories)^72^, with an exponential prior with mean = 0.5 on the alpha parameter; a Normal prior on the tree root height (in years before present), with μ = 6,912 and σ = 1,400; a Normal prior on the most recent common ancestor (MRCA, in years before present) of all CTVT samples, with μ = 1,900 and σ = 100; a strict clock, with an exponential prior with mean 6.87 x 10^−7^ substitutions per site, per year on the clock rate, as estimated in Baez-Ortega *et al.* (2019)^57^. CTVT samples were constrained to be monophyletic. Inferences were based on 20,000,000 iterations of MCMC, with the first 2,000,000 (10%) discarded as burn-in. BEAST results are available in Supplementary Data 4).

### Mutation density

This describes the analysis presented in Figure 1D and Supplementary Figure 3. For Figure 1D, we selected tumours 3131T and 851T to be representative of CTVT-A and CTVT-B–G, respectively; in Supplementary Figure 3, all tumours were included. We counted the number of mutations occurring in tumours in non-overlapping 10 kilobase windows in regions intersecting the segments of N-HT1 (Supplementary Table 3). For each segment we also counted the number of mutations in 10 kilobase windows drawn from genomic regions immediately flanking N-HT1. Mutation counts from each bin were normalised by copy number. For each N-HT1 segment we compared the mutation density in an equal number of windows from the two categories (intersecting N-HT1; flanking N-HT1).

### Time of N-HT1 origin

This describes the analysis presented in Figure 3A. We examined copy number segments in genomic loci intersecting N-HT1 (Supplementary Table 3) and determined the most frequently occurring copy number state in each copy number segment among CTVT-A tumours and among CTVT-B–G tumours. We selected segments in which the most frequent copy number state in CTVT-B–G was zero, one or two, and in CTVT-A was a single integer step higher than in CTVT-B–G (step-change = 1). In these segments we counted the number of C-to-T mutations occurring at CpG sites ([C>T]G) within 1 kilobase windows in all tumours.

Using CTVT-B–G tumours, we computed each bin’s mean [C>T]G per CpG site per CTVT parental chromosome copy. We subtracted this from the equivalent bin count in each CTVT-A tumour, adjusting for copy number. The mutation count that remained after this subtraction was assigned to N-HT1. We took the mean of these counts across all bins in all CTVT-A tumours, and obtained a value of 6.89×10^−4^ [C>T]G per CpG site for N-HT1. The published CTVT mutation rate^73^ was used to convert this to a time estimate (Figure 3A).

### N-HT1 haplotype inference

#### Data set

We obtained previously published SNP variant genotypes from 967 modern and ancient dogs and wild canids^32,33^. We subsetted this to 143,306 biallelic transversion variants found within genomic regions spanned by N-HT1 (Supplementary Data 1), and genotyped these in 47 CTVT tumours and their 46 matched hosts using Platypus v0.8.1 using the options `--bufferSize=10000 --minPosterior=0 --minReads=3 --getVariantsFromBAMs=0`, using the published SNP panel VCF table as a custom source file via the option `--source`. If any variants failed to be genotyped, the analysis was repeated for these positions using `--minReads=0` and --`outputRefCalls=1`.

The N-HT1 genotype was inferred by subtracting allele counts belonging to CTVT parental chromosomes (inferred using CTVT-B–G) from allele counts observed in CTVT-A using the method outlined below.

#### Selection of genomic regions with a copy number step change

First, we extracted genomic copy number for each tumour at each of the 143,306 SNPs (copy number data are available in Supplementary Data 3). Each SNP was assigned a “CTVT-A mode copy number” (the most frequent copy number state at that position among CTVT-A tumours) and a “CTVT-B–G mode copy number” (the most frequent copy number state at that position among CTVT-B–G tumours). Next, we identified SNPs for which the argument [“CTVT-A mode copy number state” - “CTVT-B–G mode copy number state” = 1] was true. This identified the set of SNPs occurring in intervals for which the difference in mode copy number state between CTVT-A and CTVT-B–G was exactly 1. Only these 124,209 SNPs were included in subsequent analyses.

#### Estimation of allele counts

Next, at each SNP position in each tumour we made an estimate of the copy number of reference and alternative alleles, based on total copy number and purity-corrected variant allele fraction (VAF). First, at each SNP position we selected tumours whose copy number at the selected position matched the mode copy number for its group (CTVT-A or CTVT-B–G). We then used the thresholds described in the table below to assign each SNP in each tumour an alternative allele copy number (n_Alt_), i.e. the number of alternative allele copies present. In this table, the stated VAF threshold designates the boundary between states for alternative allele copy number estimation. For example, for total copy number 2, VAF less than 0.25 implies n_Alt_ = 0; VAF of 0.25 or greater but less than 0.75 implies n_Alt_ = 1; and VAF of 0.75 or greater implies n_Alt_ = 2. Thresholds were determined by inspection of VAF density at each copy number state.

| Total copy number | VAF thresholds | Alternative allele copy number ($n_{Alt}$) |
| --- | --- | --- |
| 0 | N/A | 0 |
| 1 | 0.5 | 0, 1 |
| 2 | 0.25, 0.75 | 0, 1, 2 |
| 3 | 0.15, 0.5, 0.82 | 0, 1, 2, 3 |
| 4 | 0.12, 0.39, 0.64, 0.86 | 0, 1, 2, 3, 4 |

The number of reference alleles at each position (n_Ref_) was determined by subtracting n_Alt_ from the mode total copy number assigned to that SNP position (recall that at each SNP position only tumours whose copy number at the selected position matched the mode copy number for its group (CTVT-A or CTVT-B–G) were included in this analysis). Each variant was then tagged in each tumour with an allele count status, i.e. number of reference alleles (n_Ref_) and number of alternative alleles (n_Alt_).

#### Inference of CTVT genotype

At each variant position falling within the regions identified above, in which difference in mode copy number state between CTVT-A and CTVT-B–G was exactly 1, we took all CTVT-B– G tumours for which the sample copy number state matched the mode copy number state. Using only these tumours, we selected the mode n_Ref_ and mode n_Alt_ for each SNP. These were retained and used subsequently during inference of the N-HT1 genotype.

In addition, genotypes were collapsed and inferred to be one of 6 ancestral states as follows.

| CTVT ancestral state | Definition |
| --- | --- |
| 0/0 | Copy number $\geq 2$, $n_{\text{Alt}} = 0$; $n_{\text{Ref}} \geq 2$ (only Ref alleles present) |
| 0/1 | Copy number $\geq 2$, $n_{\text{Alt}} \geq 1$; $n_{\text{Ref}} \geq 1$ (Ref and Alt alleles present) |
| 1/1 | Copy number $\geq 2$, $n_{\text{Alt}} \geq 2$; $n_{\text{Ref}} = 0$ (only Alt alleles present) |
| 0 | Copy number 1, $n_{\text{Alt}} = 0$; $n_{\text{Ref}} = 1$ (Ref allele present) |
| 1 | Copy number 1, $n_{\text{Alt}} = 1$; $n_{\text{Ref}} = 0$ (Alt allele present) |
| ./. | Copy number 0, $n_{\text{Alt}} = 0$; $n_{\text{Ref}} = 0$ (No alleles inferred) |

These states represent the inferred genotype of the most recent common ancestor of CTVT-A and CTVT-B–G and were used as the CTVT genotype in population genetics analyses outlined below.

#### Inference of N-HT1 genotype

In the same regions with mode copy number step change of 1 described above, at each SNP position we selected CTVT-A tumours whose sample copy number state matched the CTVT-A mode copy number state. At each SNP position in tumours that fulfilled this criterion, we identified the CTVT-A mode n_Ref_ and mode n_Alt_.

The N-HT1 haplotype was inferred, at each SNP position, as follows.

N-HT1 n_Ref_ = [CTVT-A mode n_Ref_] - [CTVT-B–G mode n_Ref_]

N-HT1 n_Alt_ = [CTVT-A mode n_Alt_] - [CTVT-B–G mode n_Alt_]

Sites at which N-HT1 n_Ref_ + n_Alt_ ≠ 1 were discarded, and these sites were not included in subsequent analyses.

Genotypes of N-HT1 and CTVT across 124,209 sites inferred using this method are available in Supplementary Data 1.

### Validation of N-HT1 genotype and phasing mutations using long read DNA sequencing

We identified 120,870 variants (somatic and germline variants, indels and single nucleotide variants) within the N-HT1 region from our Somatypus variant set. We repeated the ancestral reconstruction of the genotypes of N-HT1 and CTVT at 69,105 germline SNP positions.

Sample 2169Ta-Dog (CTVT-A) PacBio HiFi long DNA sequence reads were genotyped at all 120,870 variant sites. Reads that had zero mismatches compared to the inferred ancestral N-HT1 genotype, and also had at least one mismatch to any inferred ancestral CTVT homozygous positions, were designated as originating from N-HT1. Conversely, any reads with zero mismatches to CTVT and at least one mismatch with N-HT1 were designated as originating from CTVT. Any mutations that co-occur on N-HT1 reads were considered to be phased to N-HT1; mutations co-occurring on CTVT reads were considered phased to CTVT.

### Population genetics

#### The genotyped panel

We merged the inferred genotypes of CTVT and N-HT1 across the region covered by N-HT1 with the panel of SNP variant genotypes from 967 modern and ancient dogs and wild canids^32,33^ described above. The original set of samples in the germline panel were filtered to remove technical replicates, samples marked as outliers by authors of the original studies^32,33^ and two samples that were offspring of other samples present in the data set. The resulting matrix comprised 893 samples genotyped across 124,209 sites. Haploid positions (regions of copy number 1 in CTVT or the N-HT1 haplotype) were inputted as homozygous for the single inferred allele, and regions of copy number 0 in CTVT were inputted as missing data. Published ancient DNA genotypes were used^32,33^. The data was restricted to 124,209 transversion variants in order to avoid biases introduced by DNA degradation artefacts in ancient DNA samples^32,33^. This matrix, which is available in Supplementary Data 4, was used as input for the analyses described below.

#### Linkage pruning

Principal component and ADMIXTURE^24^ analyses require that SNPs in linkage disequilibrium be pruned prior to the analysis. This was done using Plink v1.9^25^, using the commands

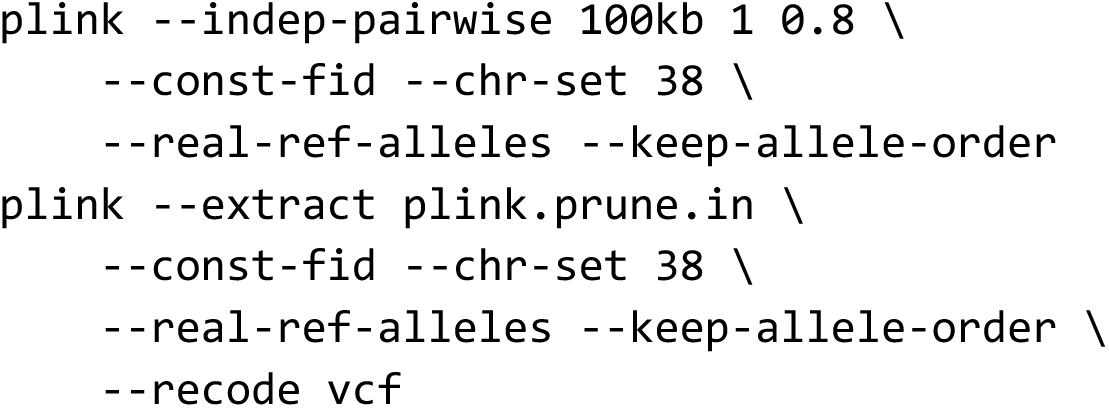

54,873 variants remained after pruning.

#### Principal Component Analysis

Principal component analysis (PCA) was done using smartpca version 18140 from

EIGENSOFT version 8.0.0^26,27^, using the following parameters:

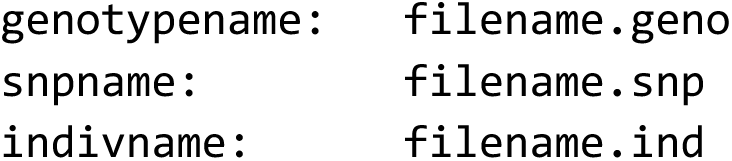
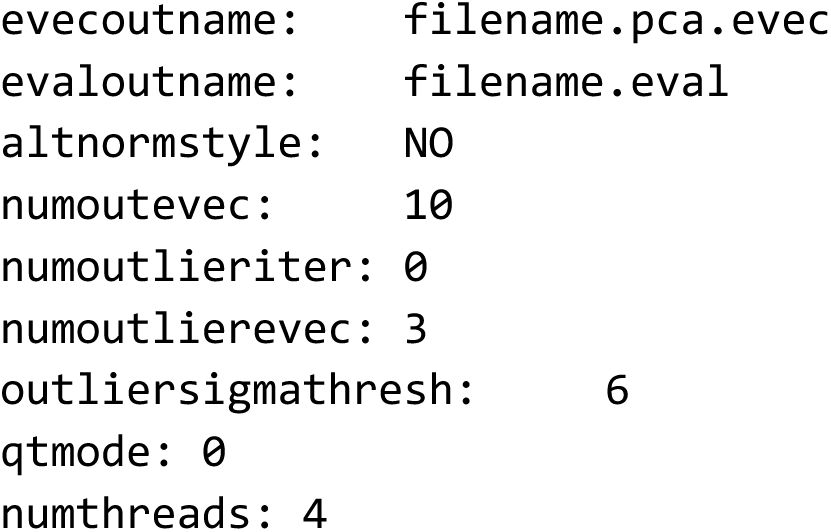

File format conversion from VCF to PED format was done using Plink v1.9^25^, and from PED to EIGENSTRAT using `convertf` from `EIGENSOFT` version 8.0.0^26,27^. The Plink-generated PED file was edited to replace ‘-9’, used by Plink to represent missing populations, with ‘3’, which is expected by `convertf` and `smartpca`.

We used a reduced set of 713 samples for PCA. Wild canid samples were not included, as initial results showed that the first principal component strongly separated this outgroup from the ingroup. Additionally, dogs of unknown or mixed breeds were removed. Populations used to build the PCA, and those plotted in Figure 3E, are listed in Supplementary Table 8.

### *f_4_-*statistics

*f_4_* statistics were calculated using the R package admixtools v2.0.0^30^. We calculated *f_4_* using Andean fox as population 1, either N-HT1 or CTVT as population 2, German shepherd as population 3, and each remaining population in turn as population 4. Block jackknifing was used to estimate the standard error in each *f_4_* statistic. We used `admixtools::f4` with parameters `auto_only=FALSE`, `blgsize=50000`, and `allsnps=TRUE`. Populations used are listed in Supplementary Table 8.

### Admixture analysis by model-based clustering

We used the ADMIXTURE v1.3.0 software^24^ to infer streams of individual ancestries in the populations represented in our data. For simplicity we inferred three latent ancestors. SNP data overlapping N-HT1 was linkage pruned as described above. The same subset of populations as was used for *f_4_* statistic was used. Because ADMIXTURE works on individual samples, for populations made up of multiple individuals we pooled the individual results obtained from ADMIXTURE as a post-processing step. Populations used are listed in Supplementary Table 8. Plotting code was adapted from https://github.com/mishaploid/Bo-demography/blob/86e618b215/src/plot_admixture_results.R.

### Mutational Signatures

Mutations occurring within the N-HT1 footprint were categorised as “trunk”, phased to CTVT parental chromosomes and occurring before the divergence between CTVT-A and CTVT-B–G; “CTVT-A”, phased to CTVT parental chromosome and occurring post-divergence in the CTVT-A lineage; and “N-HT1”, phased to N-HT1. They were identified using information from the Somatypus variant calls and the PacBio HiFi reads as follows.

**<u>Somatypus calls (all samples)</u>**

1. Select all mutations present in any CTVT-A sample – these are CTVT-A + Trunk mutations (n=17025)
2. Select all mutations present in any CTVT-B–G sample – these are CTVT-B–G + Trunk mutations (n=23243)
3. Find the intersection of the two groups – these are trunk-only mutations (n=10381)
4. Remove trunk-only mutations from (1.) – these are CTVT-A-only mutations (n=6644)
5. Remove trunk-only mutations from (2.) – these are CTVT-B–G-only mutations (n=12862)

**<u>PacBio reads (2169Tb only)</u>**

1. Select all mutations from N-HT1-identified reads – these are N-HT1 specific mutations (n=955)
2. Select all mutations from CTVT-identified reads – these are CTVT-A + Trunk mutations (n=10314)
3. Find any mutations common to both groups (6.) and (7.) – these are errors to filter out (n=39)
4. Remove errors from group (6.) – these are N-HT1 specific mutations (n=916)
5. Remove errors from group (7.) – these are CTVT-A + Trunk mutations (n=10275)

**<u>Combined selection (2169Tb only)</u>**

- Trunk = intersection of (3.) and (10.) (n=8339)
- CTVT-A = intersection of (4.) and (10.) (n=1927)
- N-HT1 = intersection of (4.) and (9.) (n=903)

Mutation spectra corresponding to these groups were plotted using sigfit^31,32^ (Figure 3B, Supplementary Figure 4).

### Mitochondrial phylogenetic tree

Mitochondrial variants were called from 768 publicly available CTVT tumours (n=390) and matched hosts (n=378)^32,33^, as well as 31 publicly available ancient DNA samples^32,33^, using the Somatypus v1.3 variant calling pipeline^69^, modified for use on the mitochondrial genome. These modifications were: disable the filter that removes single base substitution variants with median read coverage less than 20 and median VAF less than 0.2 or median VAF greater than 0.9 that are within 200bp of an indel; disable the filter that removes all single base substitution variants that have VAF greater than 0.9 in all samples; and pass additional arguments to Platypus v0.8.1^52^

--rmsmqThreshold=20

--qdThreshold=5

--maxReads=50000000

--bufferSize=2500

Single base substitution variants in the hypervariable region (positions 16110-16450) were excluded. Otherwise, single base substitutions present with a variant allele read depth of 3 or higher and a variant allele fraction (VAF) greater than 0.5 were used to construct a phylogenetic tree using the software IQ-TREE v2.2.5^71^, with substitution model GTR+G{4}^45^. Tumour specific variants were identified by comparison with the matched host. Variants occurring in both the tumour and the host have a VAF of 1. Variants occurring in the tumour and absent from the matched host occupy a purity-dependent VAF band, allowing selection of tumour-specific variants in samples of any purity (although samples with apparent purity below 5% were excluded). Node support was evaluated using 1000 ultrafast bootstrap replicates^48^. *Canis latrans* mitochondrial reference sequence DQ480510.1 was used as an outgroup to root the tree. Data are available in Supplementary Data 5.

### Gene expression

#### Identification of genes on N-HT1

Gene locations for the CanFam3.1 assembly^72^ were downloaded from Ensembl BioMart^49^, release 104. Protein coding genes were associated with the loci spanned by N-HT1 if their start or end positions intersected the coordinates of N-HT1 (Supplementary Table 4). Genes were considered truncated if they intersected N-HT1 but extended beyond its segment boundaries.

#### Transcript variant genotyping

Allele specific read counts at 775 variant positions occurring within exons in genes covered by N-HT1 were obtained from the RNAseq data using alleleCounter v2.1.2 (https://github.com/cancerit/alleleCount). Reads marked as PCR duplicates were removed. Expression counts were normalised over samples using size factors computed with DESeq2^73^.

#### Relative gene expression in the host, tumour and N-HT1

Normalised expression counts were collected for all single nucleotide variants (including germline SNPs and somatic mutations) spanned by N-HT1, for each RNAseq sample. Variants were annotated with their genotype in CTVT, N-HT1 and matched host using Illumina short read and PacBio long read phasing information.

We further annotated the table with the “informative” status of each variant. An allele was informative about relative expression levels between the three sources (host (corresponding to gene expression from tumour-infiltrating host cells), CTVT and N-HT1) if the expressed allele occurs uniquely in one of these sources. For example, for the alleles A and B, if the genotypes of both the host and CTVT are AA, and the genotype of N-HT1 is B, then the allele B is informative about the expression level of N-HT1. Similarly, if both N-HT1 and CTVT are BB, while the host is AB, then the allele A is informative about the expression of the host. If N-HT1 is absent, CTVT is AA and the host is BB, then allele A is informative about CTVT and allele B is informative about the host. We annotated the informative status of each variant according to the exhaustive combinations of these grouped genotypes. These data are provided in Supplementary Table 5. “A” refers to reference allele, “B” to alternative allele.

Normalised expression was divided by the copy number of the informative allele in the relevant group to obtain a normalised estimate of expression per genomic copy. Data points which were informative on the same gene in the same category (tumour-infiltrating host, CTVT, N-HT1) were pooled across tumours, and the mean of these for each of the 73 genes carrying N-HT1-informative alleles is presented in Figure 4B. All 73 genes also carried variants informative of tumour-infiltrating host expression, and 71 carried variants informative of CTVT expression. 95% confidence intervals were constructed using the standard error of the mean. We quantified the relationships between N-HT1 and CTVT expression, and N-HT1 and host expression, using linear regression, using the `lm` function implemented in R^57^.

### Cytogenetics

#### Metaphase preparation

3mm^3^ CTVT tumour biopsies were finely minced in calcium- and magnesium-free phosphate buffered saline (PBS) (Sigma-Aldrich, St Louis, USA). The resulting cell suspension was filtered through a 100μm cell strainer and centrifuged at 1000rpm for 5 minutes before being resuspended in 10ml Dulbecco’s modified eagle medium (DMEM) (Sigma-Aldrich, St Louis, USA) with 10% foetal bovine serum (Sigma-Aldrich, St Louis, USA), 1% penicillin/streptomycin (Thermo Fisher Scientific, Waltham, USA) and 1μg Colcemid (Thermo Fisher Scientific, Waltham, USA). Cells were then incubated at 37°C, for 2-3 hours. Cells were then centrifuged (1000rpm, 5 minutes) and resuspended in 5ml of prewarmed (to 37°C) 0.075M potassium chloride (KCl) solution. This was followed by incubated for 10 minutes at 37°C with 30 seconds of mixing halfway through. The cells were then centrifuged again (600rpm, 5 minutes) before 5ml of a freshly prepared fixative of 3:1 pure methanol : glacial acetic acid was added drop by drop with careful agitation to prevent clumping.

Metaphase spreads from the MDCK canine cell line were prepared in the same way, except the starting material was a near-confluent flask of cells, and cells were harvested using trypsin and mechanical scraping.

#### Bacterial artificial chromosome probe preparation

Dog BAC clones CH82-448A7, CH82-283H6 and CH82-276P21 were purchased from BACPAC Genomics (Emeryville, USA). Clones were obtained as stab cultures in LB agar containing 12.5µg/mL chloramphenicol. These were streaked to single colonies, which were grown overnight at 37°C in LB medium containing 12.5µg/mL chloramphenicol. DNA extraction was performed using the QiaPREP Spin Miniprep Kit (Qiagen, Hilden, Germany). Clone identity was validated with PCR, and their genomic mapping locations in CanFam3.1 are listed below. Expected copy number for each probe in CTVT-A and CTVT-B–G is shown below, based on genomic copy number. Figure 2A shows a schematic diagram illustrating probe mapping locations on N-HT1.

| Clone | Mapping coordinates (CanFam3.1) | CTVT-A expected copy number | CTVT-B-G expected copy number |
| --- | --- | --- | --- |
| CH82-448A7 | 1:32528835-32701528 | 1 (N-HT1) | 0 |
| CH82-276P21 | 1:31017317-31189651 | 2 | 1 |
| CH82-283H6 | 7:7848088-8032891 | 3 | 2 |

### Fluorescence *in situ* hybridisation

The chromosome cell suspension was dropped onto clean slides and dried overnight. The probes were labelled with the Nick Translation Kit (Abbott Molecular, Des Plaines, USA) following the manufacturer’s instructions, with biotinylated-16-dUTP (Sigma–Aldrich, St Louis, USA), Spectrum Red-dUTP (Vysis, Abbott Molecular, Des Plaines, USA), and Spectrum Gold-dUTP (Enzo Laboratories, Farmingdale, USA). The probes were purified by precipitation, adding a 10 × excess of unlabelled canine DNA, and re-suspended in hybridization buffer (50% formamide, 10% dextran sulfate, 2 × SSC). The probes were denatured for 8 minutes at 85°C and pre-annealed at 37°C for 30 minutes.

Metaphase spread DNA was denatured in NaOH, 0.07 M, for 1 min, and the denatured probe mix was applied to the slides under a coverslip. The hybridization was carried out for two days at 37°C, in a humidified chamber. Post-hybridization washes were in 0.1 × SSC at 60°C. The biotinylated probe was detected using Streptavidin-Alexa Fluor 488 (Thermo Fisher Scientific, Waltham, USA). The slides were mounted in DAPI/Vectashield (Vector Laboratories, Newark, USA).

Images were collected with an Olympus BX-51 microscope, equipped with a JAI CVM4+ CCD camera, using Leica Cytovision Genus v7.1.

In addition to CTVT metaphases (Figure 2), FISH was performed on MDCK canine kidney cells in order to confirm expected mapping of probes.

## Data accession

Whole genome sequencing and RNA sequencing data have been deposited in ENA with accession (data deposition in progress). Additional data is available on Zenodo (https://doi.org/10.5281/zenodo.7214808). This includes:

Supplementary Data 1: Germline panel vcf (N-HT1 region)

Supplementary Data 2: Somatypus variant calls

Supplementary Data 3: CTVT copy number data

SupplementaryData 4: BEAST tree data and newick

Supplementary Data 5: Mitochondrial tree data and newick

Code relating to analyses performed in the study can be found at https://github.com/TransmissibleCancerGroup/NuclearHorizontalTransfer. Data required for running code examples is available in Supplementary Data 1–5. Ancestral inference and population genetics analyses were implemented as Nextflow pipelines^36^, available at https://github.com/TransmissibleCancerGroup/NuclearHorizontalTransfer.

## Supplementary Information

**Supplementary Table 1: CTVT sample metadata.**

Host sex: M, male; F, female. CTVT_Group refers to phylogenetic group defined in Supplementary Figure 1. The CTVT Group of tumour 3838T is not known (it belongs to one of B, C, D, E, F or G). Tasmanian devil sample metadata are available in Stammnitz *et al.* 2023^29^.

**Supplementary Table 2: N-HT1 structural variants.**

**(A)** N-HT1 structural variant features. Two softwares, Manta and SvABA, were used to identify structural variants using Illumina short reads. Breakpoint features predicted by Manta and SvABA were supported by PacBio HiFi long reads.

**(B)** PacBio HiFi long DNA sequencing reads supporting N-HT1 structural variant breakpoint junctions.

**Supplementary Table 3: N-HT1 genomic coordinates.**

Coordinates are relative to CanFam3.1.

**Supplementary Table 4: Genes encoded on N-HT1.**

Genes truncated by segment boundaries are indicated.

**Supplementary Table 5: Allele-specific gene expression data for 125 genes carrying alleles informative for CTVT, host or N-HT1**

In Figure 4B, values for ‘N-HT1 expression’ were obtained by selecting rows with value ‘ht_only’ in the column ‘informative_category’. For each gene in Figure 4B, ‘N-HT1 expression’ is given by the mean value of ‘expression’ in the rows corresponding to the relevant gene, together with its standard error. ‘CTVT expression’ and ‘Host expression’ was obtained in the same manner, selecting for ‘ctvt’ and ‘host’ in the ‘informative_category’, respectively.

**Supplementary Table 6: Annotation of non-synonymous somatic mutations occurring in *ARFGEF3*.**

Variant phasing to ‘CTVT_parental_chromosome’ or ‘N-HT1’ was performed using PacBio HiFi long sequence reads.

**Supplementary Table 7: Coordinates of homozygous deletions complemented in CTVT-A by N-HT1.**

Coordinates of homozygous deletions in CTVT-B-G in regions spanned by N-HT1.

**Supplementary Table 8: Details of samples and populations used in population genetics analyses.**

(A) Details of samples used in Figure 3E (PCA). All listed samples were used in PCA calculation, and “Plotted” indicates those plotted in Figure 3E.

(B) Details of samples and populations used in Figure 3C (F4 statistics) and Figure 3D (ADMIXTURE). Because ADMIXTURE works on individual samples, for populations made up of multiple individuals we pooled the individual results obtained from ADMIXTURE as a post-processing step.

Sample names relate to published papers^32,33^.

---

**Supplementary Figure 1:**
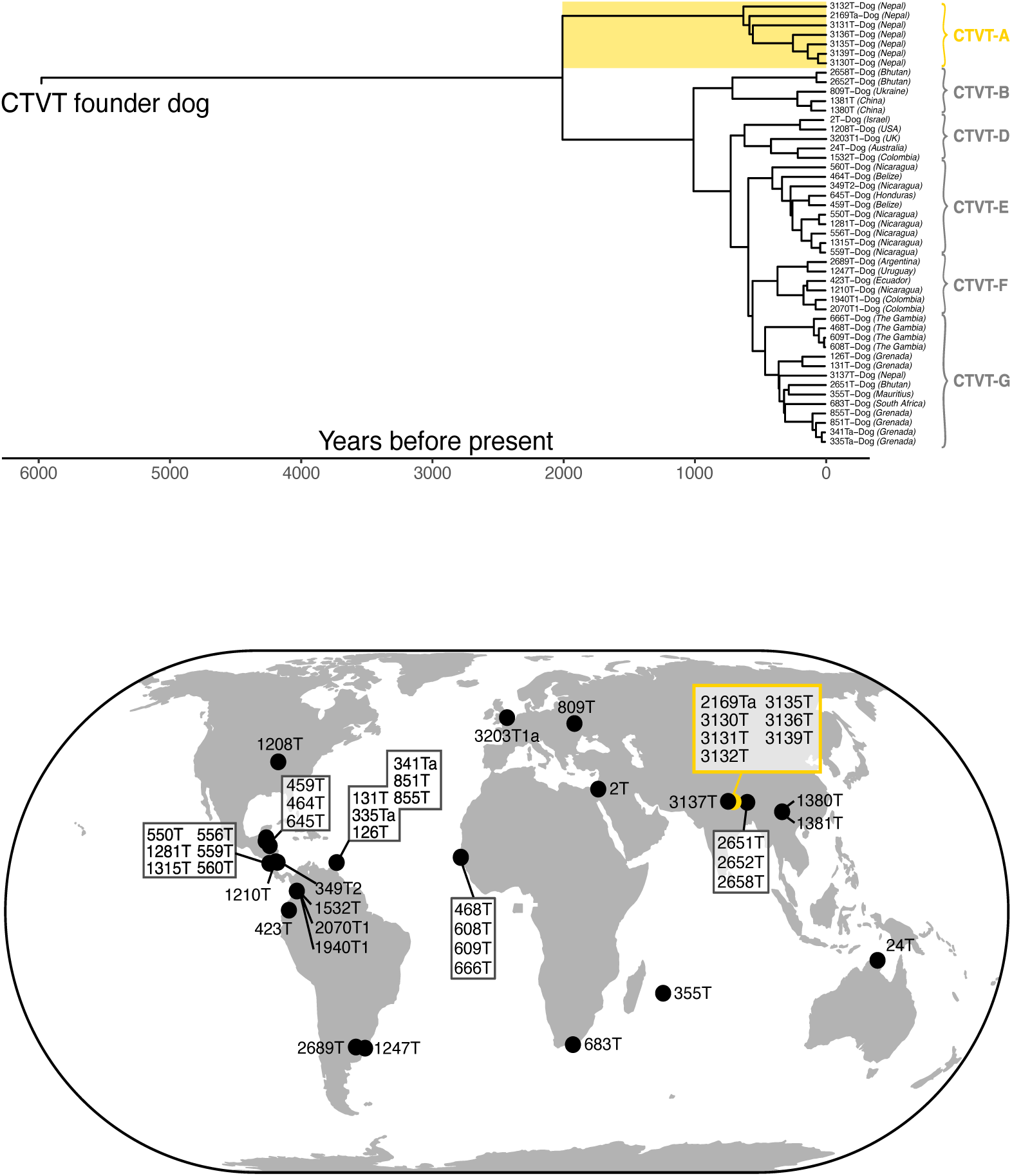
CTVT phylogenetic tree and sampling locations. Time-scaled phylogenetic tree and sampling locations of 47 CTVT tumours analysed in this study. Detailed metadata is available in Supplementary Table 1. No tumours belonging to CTVT-C, which corresponds to “Group 1 – India (Kolkata)” (Baez-Ortega *et al.* 2019) were included in the study. CTVT-A is highlighted in gold. The map is CartoDB_PositronNoLabels obtained from the R package tmap.

**Supplementary Figure 2:**
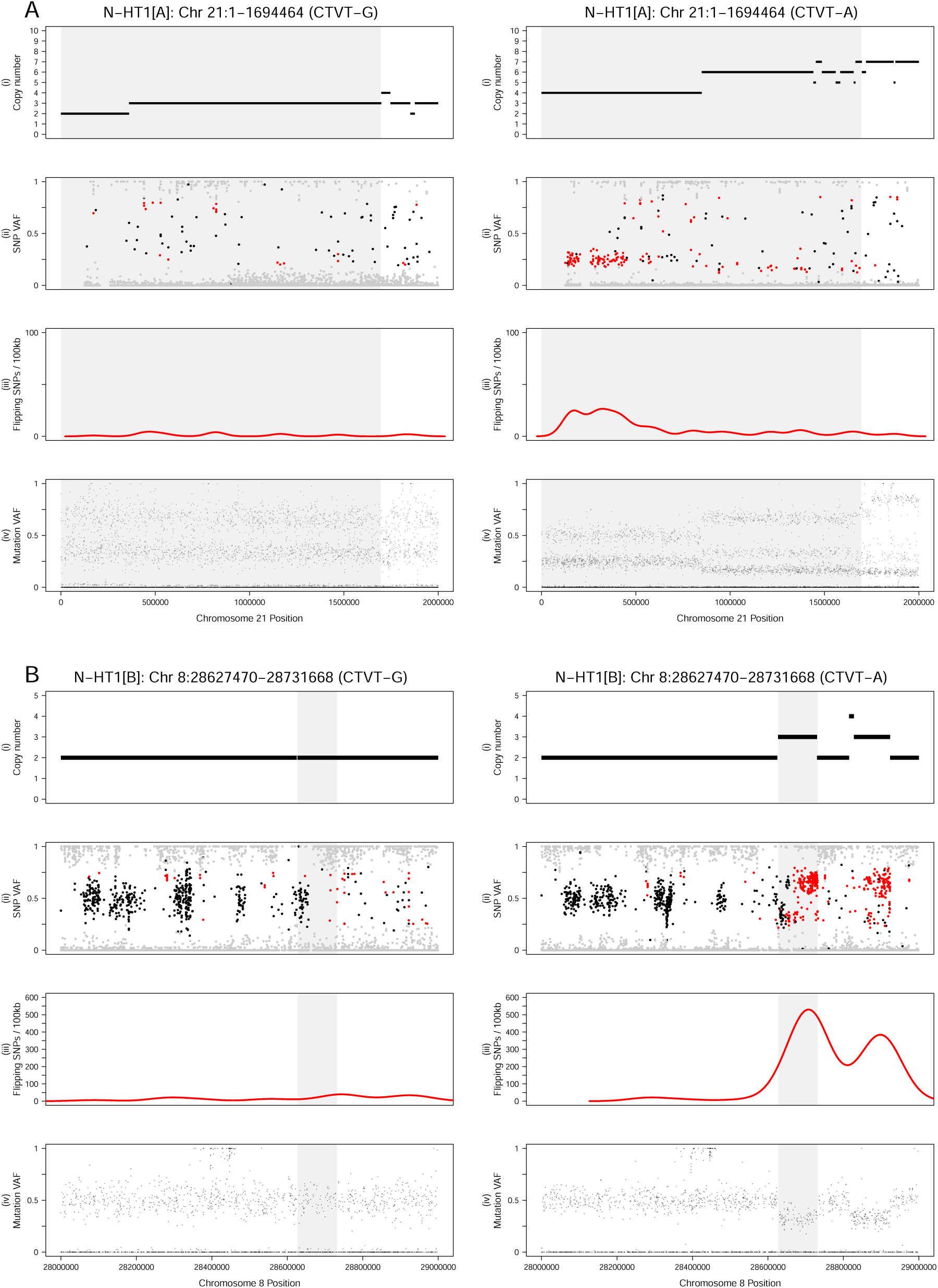
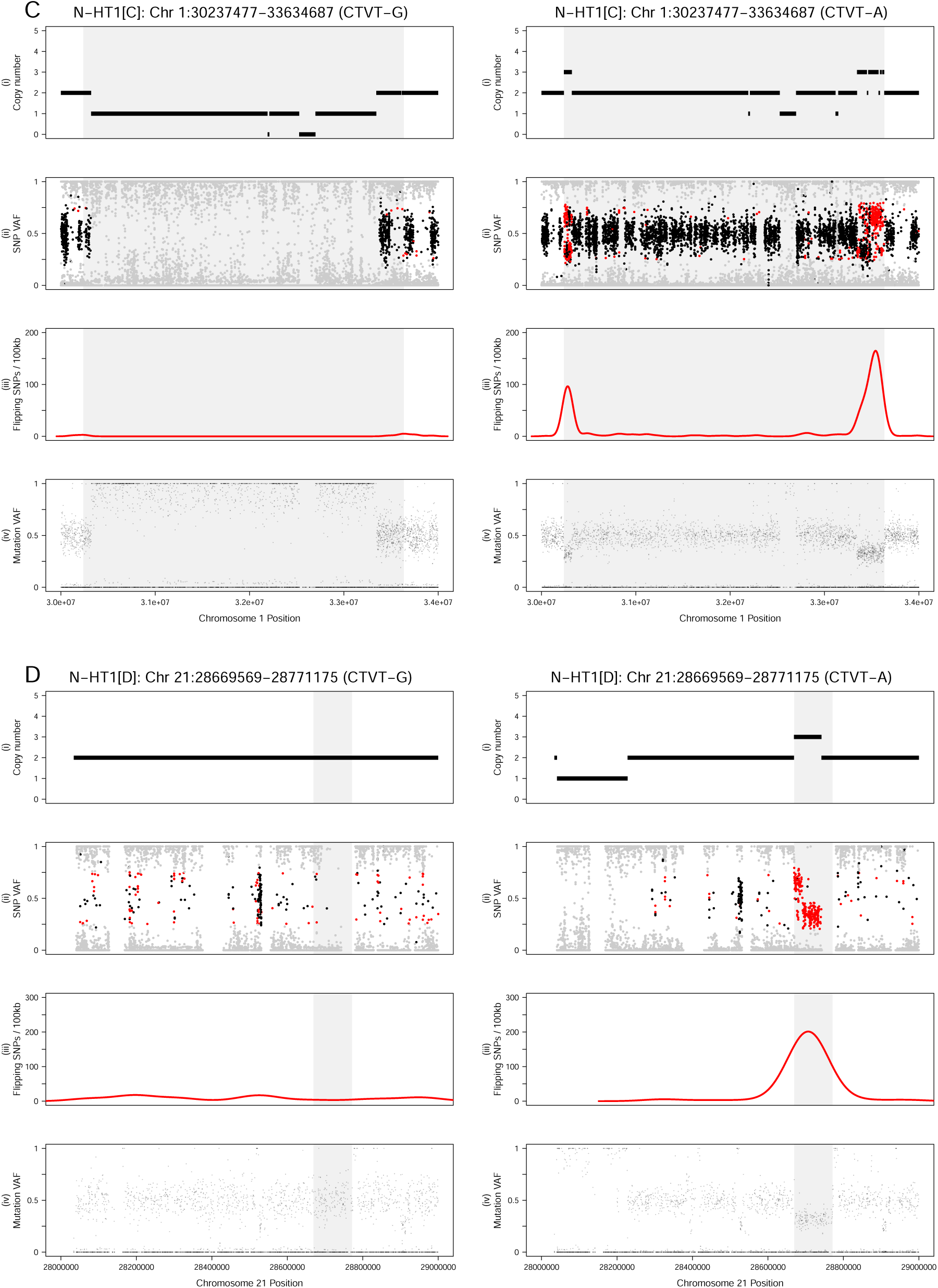
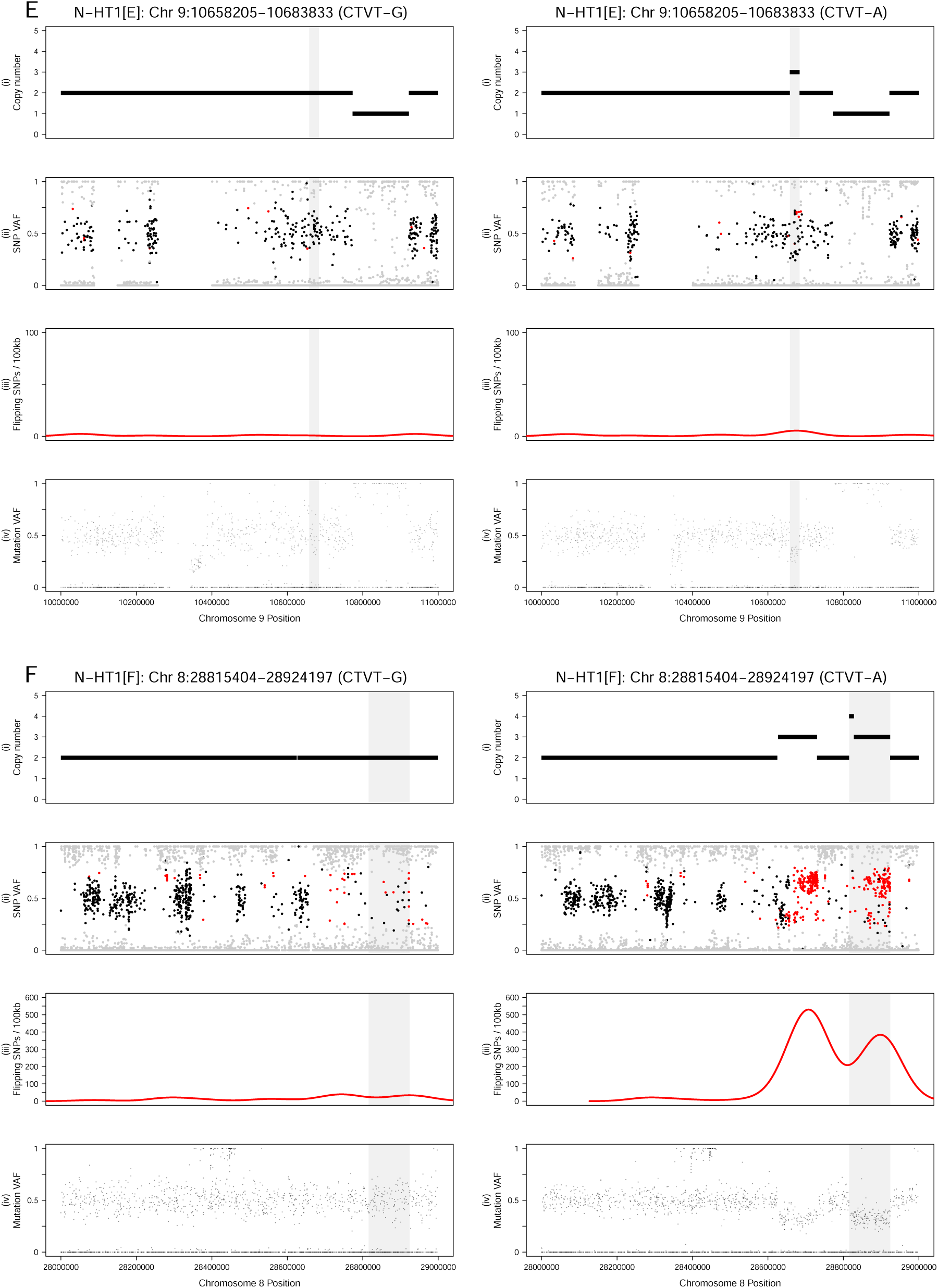
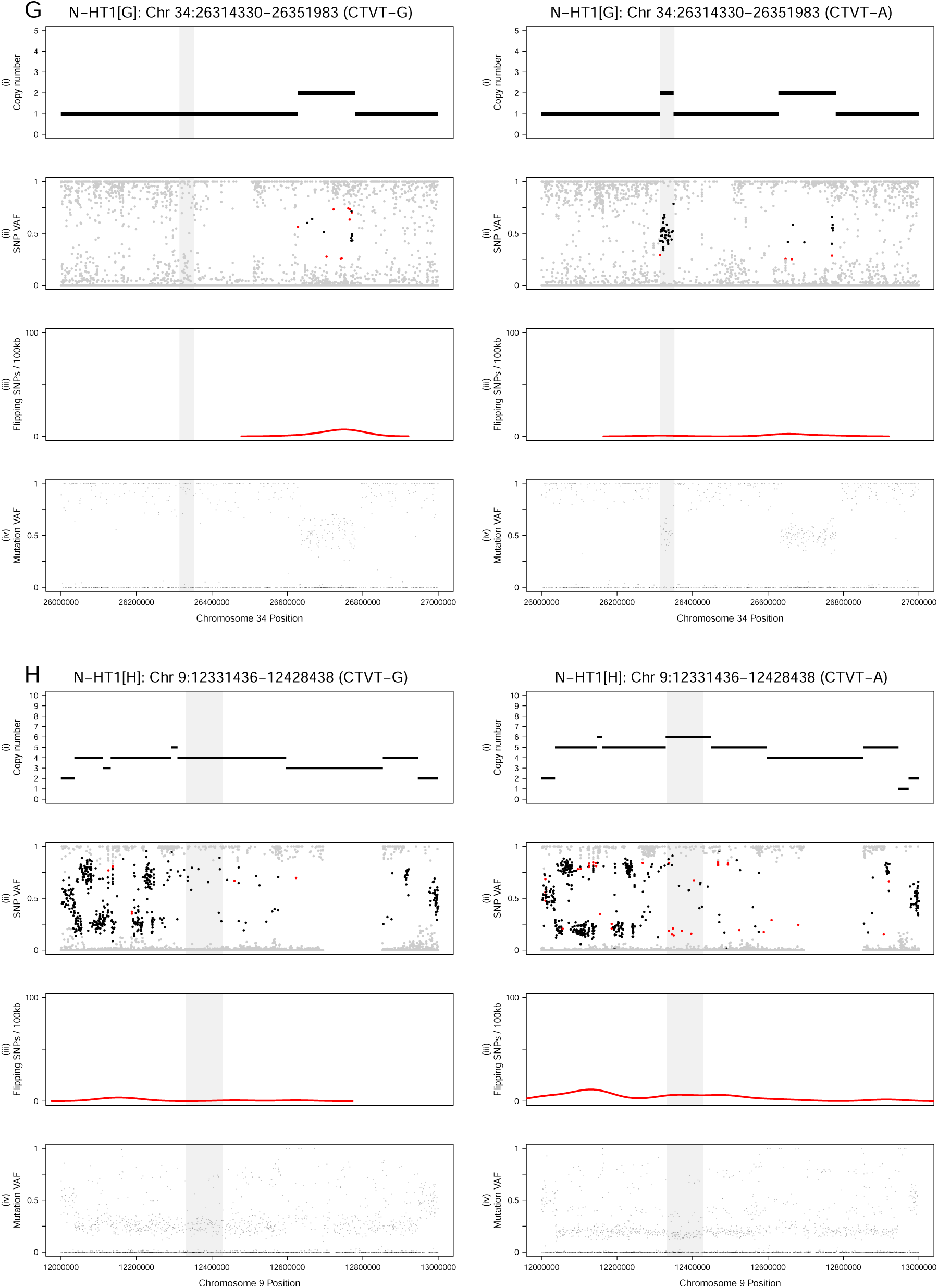
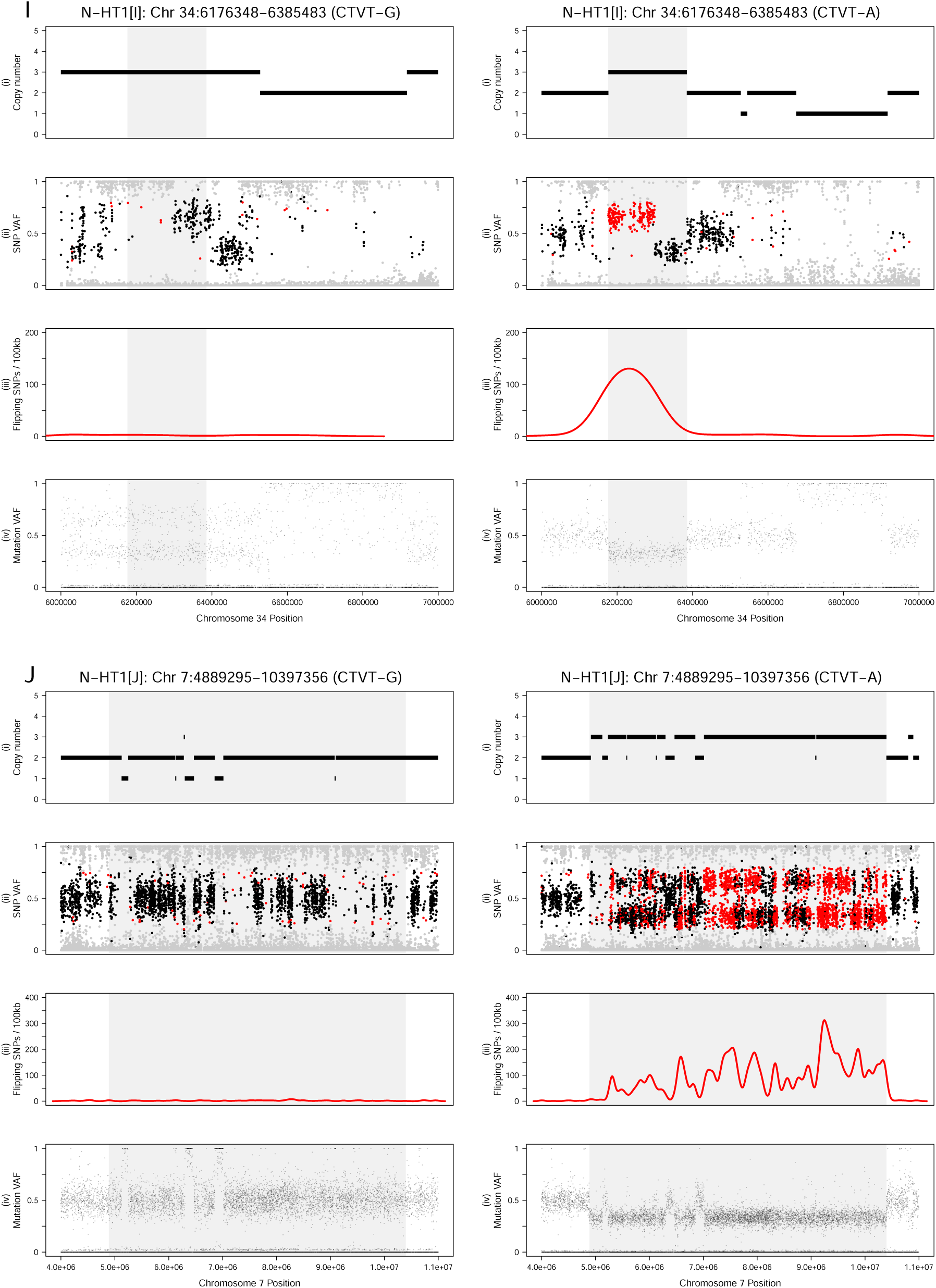
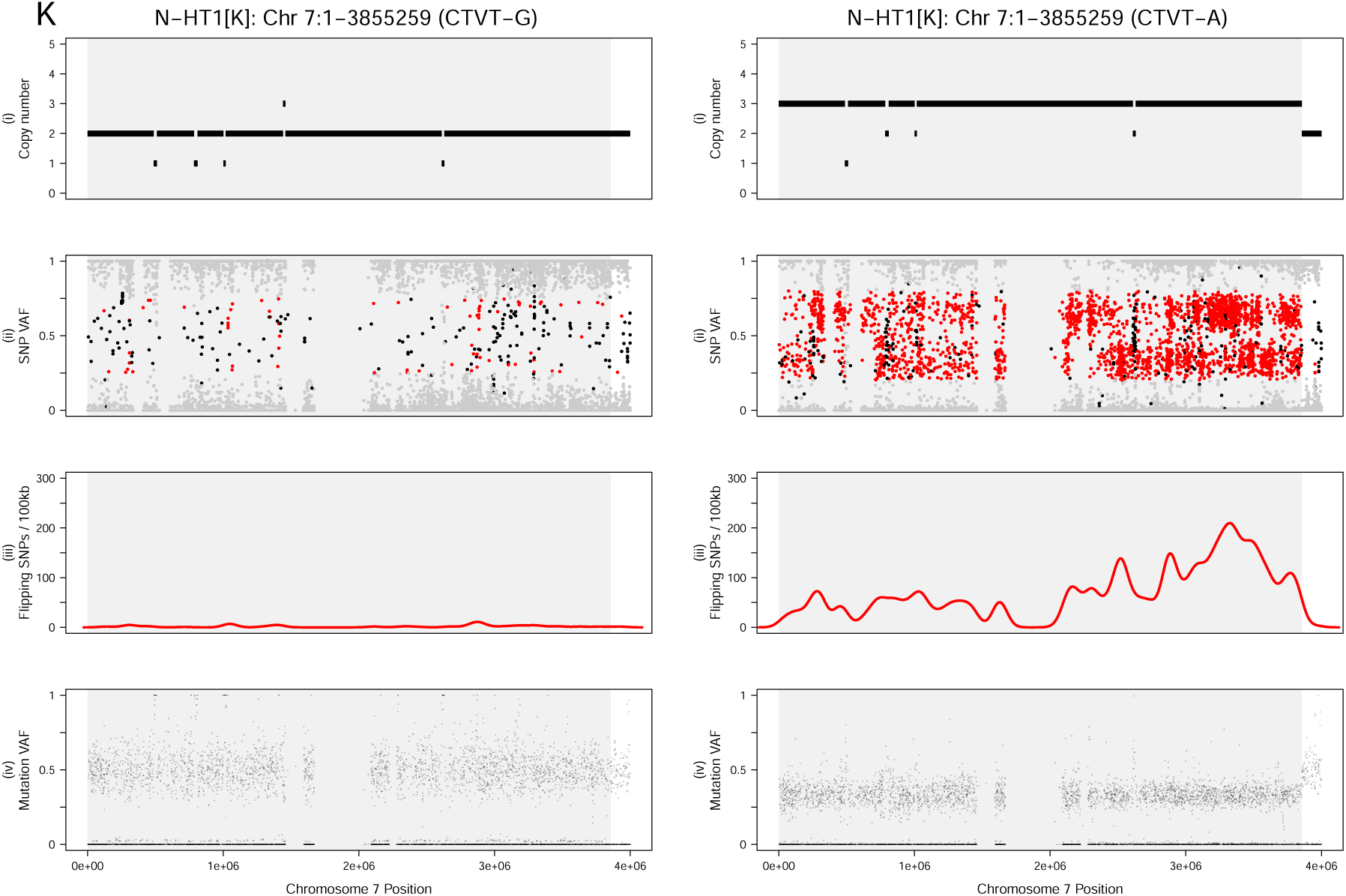
Genome data in N-HT1 segments. Data corresponding to each of the 11 segments (A–K) of N-HT1for representative tumours belonging to CTVT-B–G (left, CTVT-G tumour 851T, horizontal DNA transfer absent), and CTVT-A (right, tumour 3131T, horizontal DNA transfer present). The horizontally transferred locus is in the centre of each panel, marked with a grey background, and the identity of the N-HT1 segment (A–K, see Figure 1E) is labelled in square brackets at the top of each plot. Each set of panels show (i) copy number; (ii) SNP variant allele fraction (VAF) corrected for tumour purity; (iii) density of flipping SNPs; and (iv) somatic mutation VAF corrected for tumour purity. In (ii), SNPs with heterozygous genotype in the majority of CTVTs are coloured black; those with homozygous genotype in the majority of CTVTs but heterozygous genotype in the sample shown (“flipping SNPs”) are coloured red; all other SNPs are coloured grey. Scatterplots are limited to at most 20,000 data points, selected uniformly at random.

**Supplementary Figure 3:**
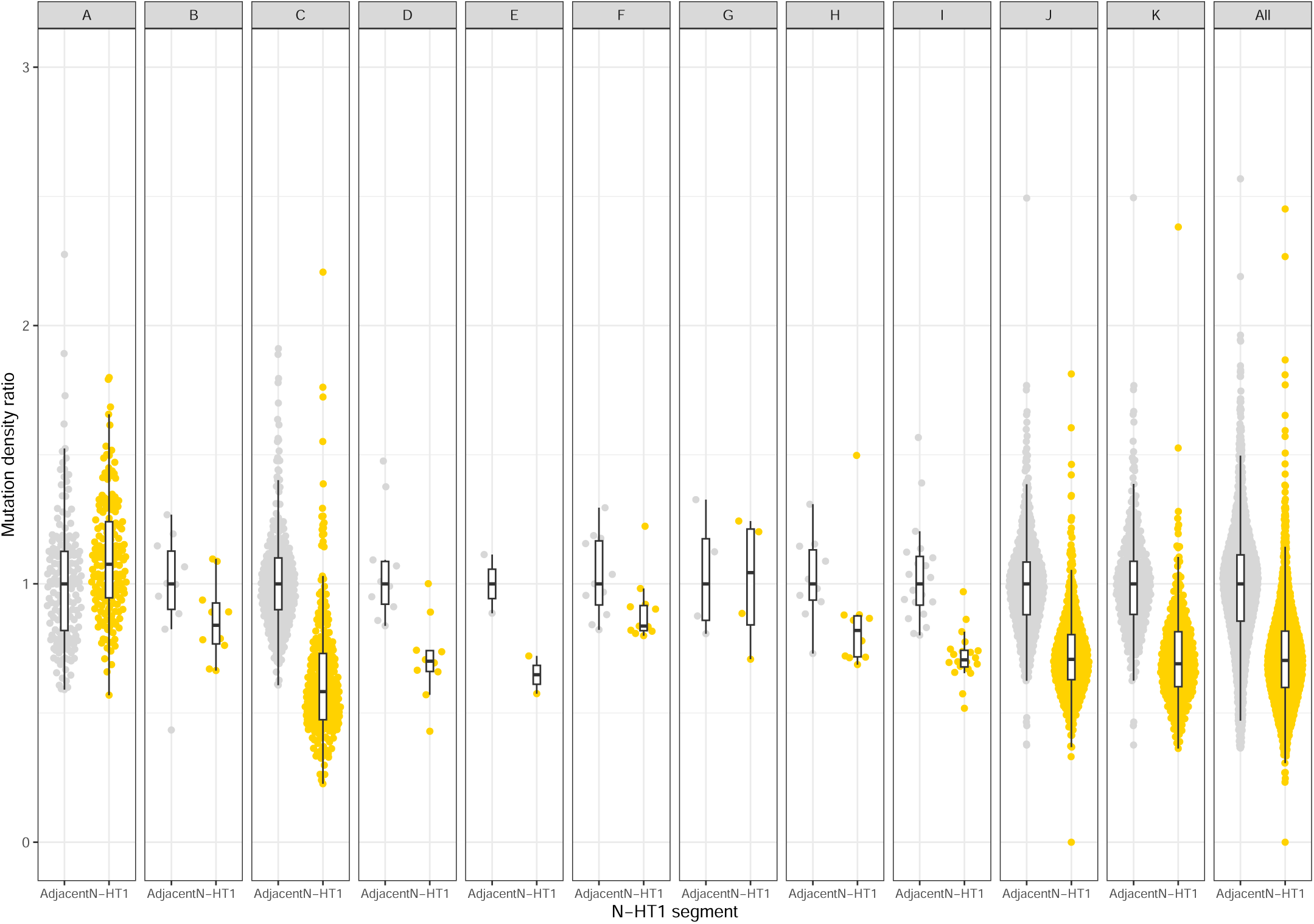
Mutation density in N-HT1 segments. Mutation density for each of the eleven segments (A–K) spanned by N-HT1. Each “N-HT1” point represents CTVT-A average mutation count per DNA copy in 10 kilobase (kb) bins, expressed as a fraction of average mutation density in the equivalent bins in CTVT-B–G tumours. “Adjacent” shows the equivalent fraction in an equally sized genomic region immediately adjacent to the designated N-HT1 segment. The median of the “Adjacent” mutation count ratio is set to 1. Decrease in mutation density specific to CTVT-A is not observed in N-HT1 segment A because additional copies of this region have been acquired in CTVT-B–G tumours through duplication events (Supplementary Figure 2 and Materials and Methods).

**Supplementary Figure 4:**
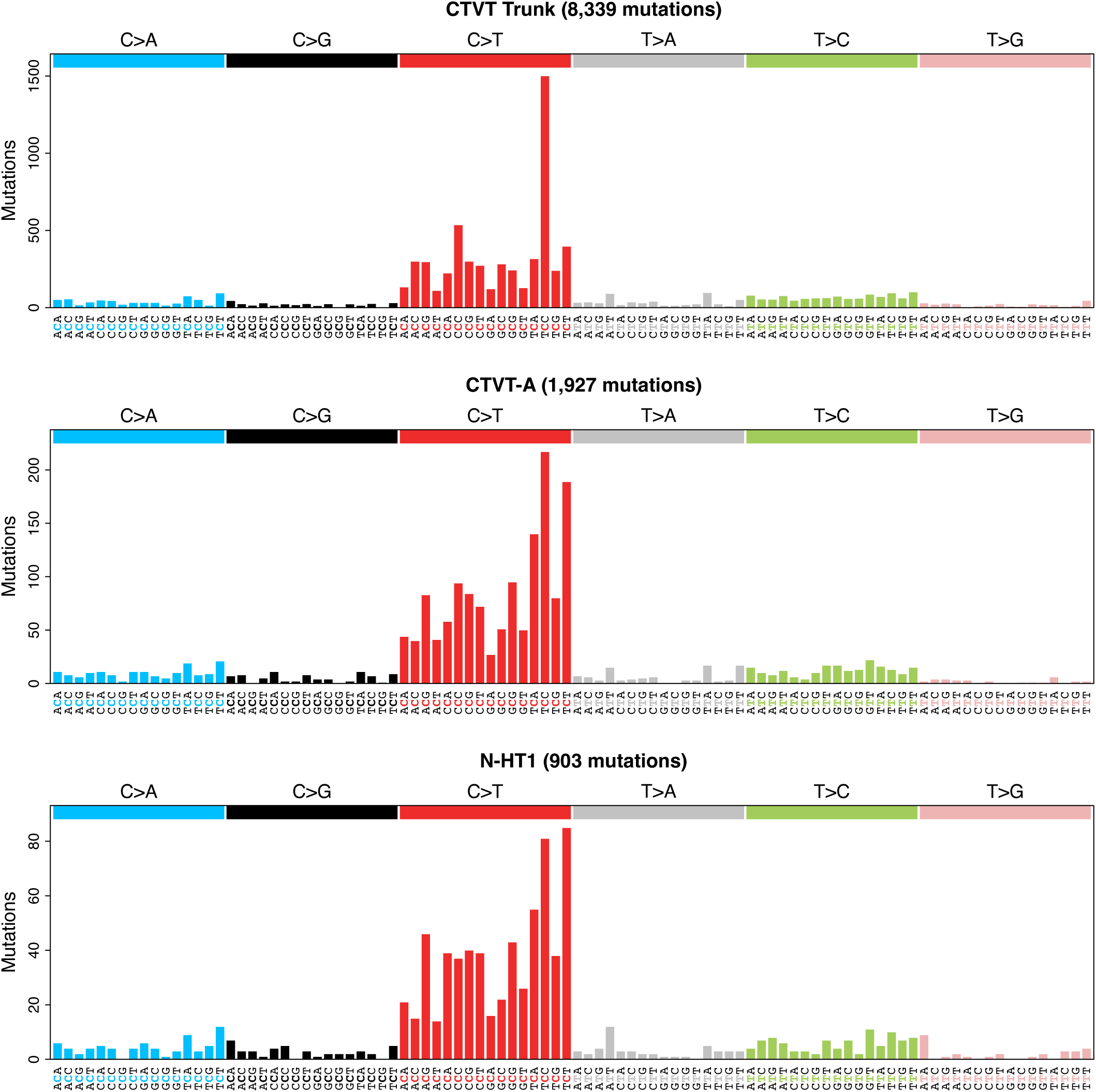
Mutation spectra. Fully annotated plots corresponding to Figure 3B.

**Supplementary Figure 5:**
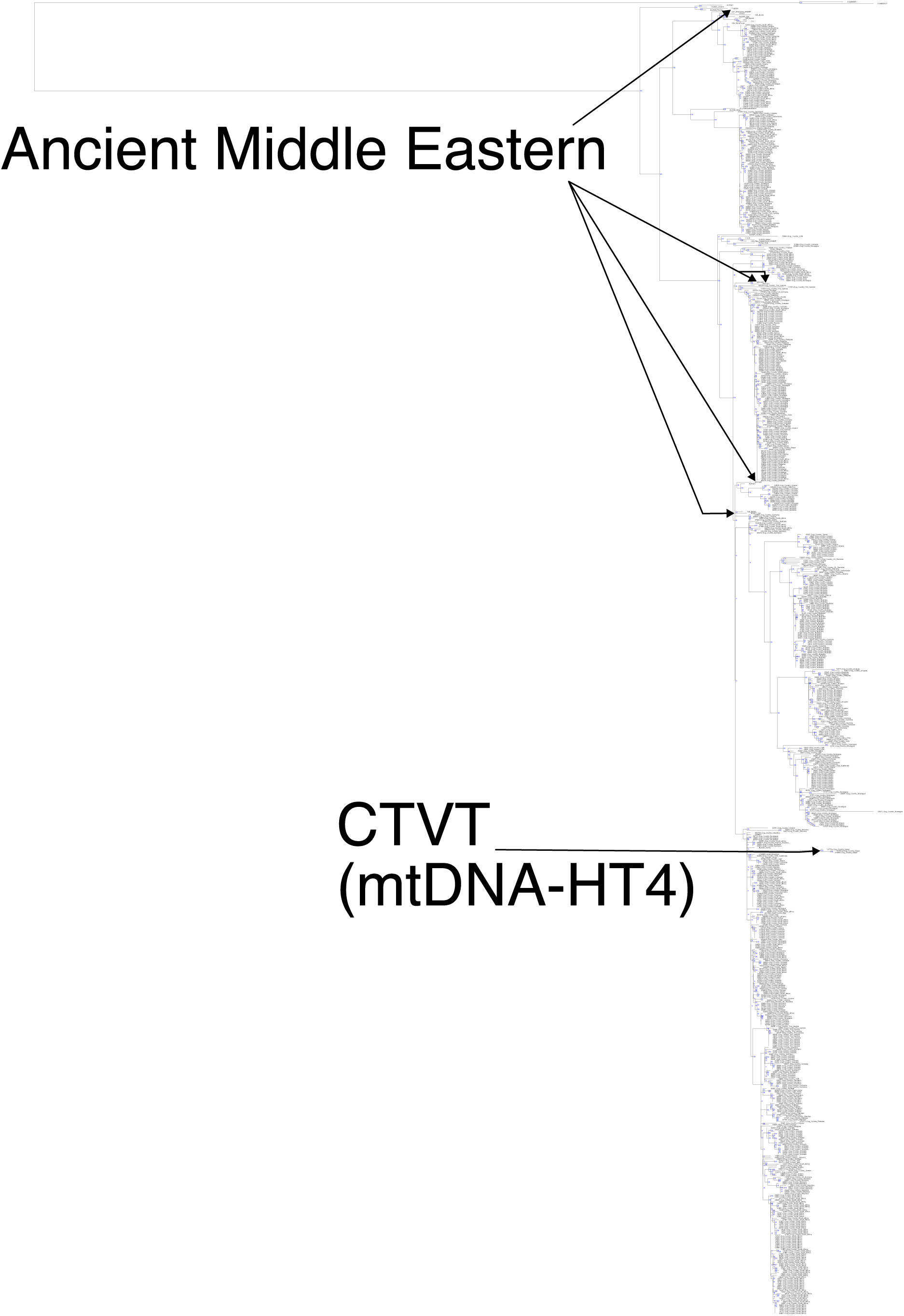
Mitochondrial phylogenetic tree. A phylogenetic tree built using maximum likelihood estimation based on 882 germline and 797 somatic mitochondrial single nucleotide variants in 390 CTVT tumours, 378 CTVT host dogs, 31 ancient dogs, and a coyote outgroup (Strakova *et al.* 2020; Bergström *et al.* 2020). CTVT-A tumours, which carry mtDNA-HT4 (Strakova *et al.* 2020), are indicated, as are mtDNAs from five ancient Middle Eastern dogs (see Figure 2C, 2D and 2E). Tumours have suffix “-T” and matched host dogs “-H”. 2174T and 2174H have had their labels switched, and “2174H” is in fact a tumour and “2174T” is in fact a host. CTVT tumours and hosts have sampling country annotated. Ancient samples are annotated with their accession identifiers (see Bergström *et al.* 2020 for more details). Bootstrap support values are based on 1000 replicates. Genotype data are available Supplementary Data 4.

## Notes

https://doi.org/10.5281/zenodo.7214808

### Competing Interest Statement

The authors have declared no competing interest.

