## Supplementary Table Legends for "Horizontal transfer of nuclear DNA in transmissible cancer"

Sample names relate to published papers^2,3^.

1.	Stammnitz, M. R. *et al.* The evolution of two transmissible cancers in Tasmanian devils. *Science* **380**, 283–293 (2023).

2.	Bergström, A. *et al.* Origins and genetic legacy of prehistoric dogs. *Science* **370**, 557–564 (2020).

3.	Bergström, A. *et al.* Grey wolf genomic history reveals a dual ancestry of dogs. *Nature* **607**, 313–320 (2022).
